# slimr: An R package for integrating data and tailor-made population genomic simulations over space and time

**DOI:** 10.1101/2021.08.05.455258

**Authors:** Russell Dinnage, Stephen D. Sarre, Richard P. Duncan, Christopher R. Dickman, Scott V. Edwards, Aaron Greenville, Glenda Wardle, Bernd Gruber

## Abstract

Software for realistically simulating complex population genomic processes is revolutionizing our understanding of evolutionary processes, and providing novel opportunities for integrating empirical data with simulations. However, the integration between simulation software and software designed for working with empirical data is currently not well developed. Here we present slimr, an R package designed to create a seamless link between standalone software SLiM 3.0, one of the most powerful population genomic simulation frameworks, and the R development environment, with its powerful data manipulation and analysis tools. We show how slimr facilitates smooth integration between genetic data, ecological data and simulation in a single environment. The package enables pipelines that begin with data reading, cleaning, and manipulation, proceed to constructing empirically-based parameters and initial conditions for simulations, then to running numerical simulations, and finally to retrieving simulation results in a format suitable for comparisons with empirical data – aided by advanced analysis and visualization tools provided by R. We demonstrate the use of slimr with an example from our own work on the landscape population genomics of desert mammals, highlighting the advantage of having a single integrated tool for both data analysis and simulation. slimr makes the powerful simulation ability of SliM 3.0 directly accessible to R users, allowing integrated simulation projects that incorporate empirical data without the need to switch between software environments. This should provide more opportunities for evolutionary biologists and ecologists to use realistic simulations to better understand the interplay between ecological and evolutionary processes.

## Introduction

Mathematical modelling and simulation are critical cornerstones of population genetic practice. At a fundamental level, empirical datasets demand analytical tool-kits that can accomodate their high complexity, and recent developments in sophisticated simulation software have the potential to provide mechanistic insight into increasingly complex evolutionary scenarios (Carvajal-Rodríguez, 2010; Haller & Messer, 2019; Hoban, 2014; Kelleher, Etheridge, & McVean, 2016; Messer, 2013; Strand, 2002; Yuan, Miller, Zhang, Herrington, & Wang, 2012). However, utilising flexible simulations requires exploration of large parameter space, which often generates large amounts of data that need sophisticated computational tools to unpack, interrogate and synthesize. Likewise, using simulations to model empirical data is an emerging field because it allows researchers to deal with complex situations where it is difficult to obtain a closed likelihood (Beaumont, Zhang, & Balding, 2002; Brehmer, Louppe, Pavez, & Cranmer, 2020; Cranmer, Brehmer, & Louppe, 2020; Marjoram, Molitor, Plagnol, & Tavare, 2003; Sisson, 2018; Torada et al., 2019; Wang et al., 2020). To facilitate more rapid and seamless interrogation and synthesis between empirical data and population genetics simulation, we present `slimr` (https://rdinnager.github.io/slimr/). `slimr` is an R package designed to link the very large and widely used ecosystem of analysis and visualization tools in the R statistical language to the SLiM scripting language (Haller & Messer, 2019), a popular, powerful and flexible population genetics simulation tool. The package creates a smooth fusion between the computational power and flexible model specification of SLiM with the advanced statistical analysis, visualisation, and metaprogramming tools of R.

### Package Description

`slimr` is an R package that interfaces with SLiM 3.0 software for forward population genetics simulations (Haller & Messer, 2019 for full details on SLiM, as well as the website at https://messerlab.org/slim/; see Messer, 2013).

`slimr` implements a Domain Specific Language (DSL) that mimics the syntax of SLiM, allowing users to write and run SLiM scripts and capture resulting simulation output, all within the R environment. Much of the syntax is identical to SLiM, but `slimr` offers additional R functions that allow users to manipulate SLiM scripts (“`slimr` verbs”) by inserting them directly into any SLiM code block. This enables R users to create SLiM scripts that explore large numbers of different parameters and also automatically produce output from SLiM for powerful downstream analysis within R.

The features of slimr fall into three categories: 1) SLiM script integrated development, 2) data input/output, and 3) SLiM script metaprogramming. The first set of features is designed to make it easy to develop SLiM scripts in an R development environment such as Rstudio, and mostly recapitulates features that SLiM users already have access to in the form of SLiMgui and QtSLiM (https://messerlab.org/slim/). The second and third features are implemented using what we call “`slimr` verbs”, allowing SLiM and R features to be combined in advanced ways. The integration between R and SliM provided by `slimr` compensates knowledgeable users of R for a lack of knowledge of SLiM, helping to lower the barrier to learning and using SLiM.

Each of the 3 categories has subcategories of features as follows:

1. Integrated Development
  a. Autocomplete and Documentation (within R) for SliM code
  b. Code highlighting and pretty printing of SLiM code
  c. Rstudio addins
  d. Run code in SLiM from R
2. Data Input/Output
  a. Automatic output generation and extraction from SLiM to R (`slimr_output()`)
  b. Insert arbitrary R objects into SLiM scripts through inlining (`slimr_inline()`)
3. Metaprogramming
  a. Code templating for SLiM scripts (`slimr_template()`)
  b. Flexible general metaprogramming tools (support for `rlang`’s `!!` and `!!!` forcing operators)

In the next section we describe each of these features in greater detail, showing examples through screenshots and code snippets.

### Integrated Development

`slimr` allows the user to write SliM code from within an R integrated development environment (IDE). `slimr` is designed to work well with Rstudio, but can be used in any R IDE. The syntax used to write SliM code is very similar to the native SliM syntax, with a few modifications to make it work with the R interpreter. As an example, here is a minimal SliM program, and its counterpart written in `slimr`.

#### SliM code

~~~
initialize()
{
  initializeMutationRate(1e-7);
  initializeMutationType(“m1”, 0.5, “f”, 0.0);
  initializeGenomicElementType(“g1”, m1, 1.0);
  initializeGenomicElement(g1, 0, 99999);
  initializeRecombinationRate(1e-8);
}
1
{
  sim.addSubpop(“p1”, 500);
}
10000
{
  sim.simulationFinished();
}
~~~

#### slimr code

The data looks like this:

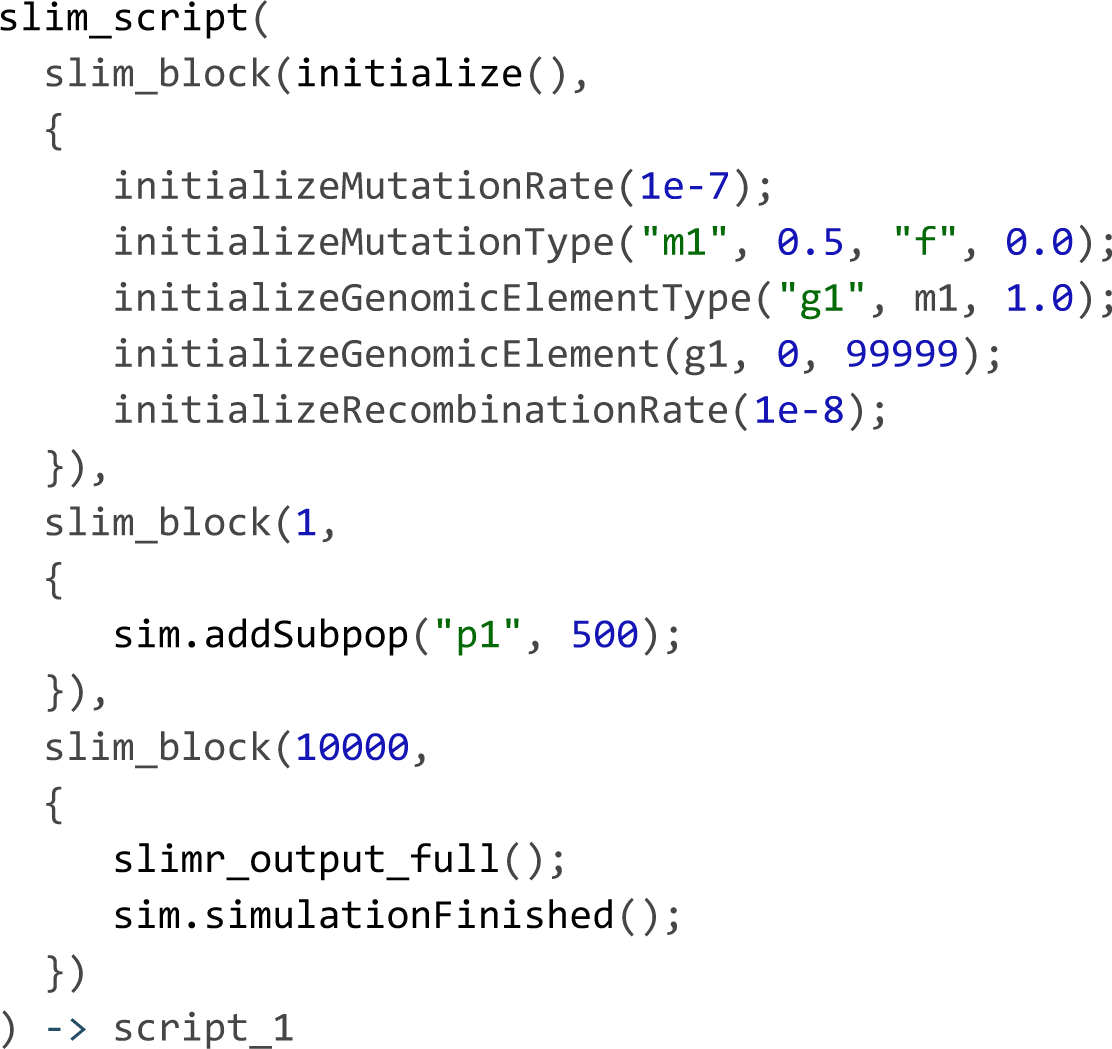

The above code assigns the script to an R object `script_1`, which can then be further manipulated, printed prettily, and sent to SliM to run. See Fig. 1 to see what the above script looks like in the Rstudio IDE, and examples of things you can do with it. A script is specified using the `slim_script` function, within which you create `slimr` coding blocks, using the `slim_block` function. The user can create as many `slimr` code blocks as desired within a `slim_script`. We’ve added a `slimr` “verb” (`slimr_output_full`), which tells SliM to output the full state of the simulation and return it to R during the execution of the block. We will discuss `slimr` verbs in more detail in the next section.

**Figure 1.**
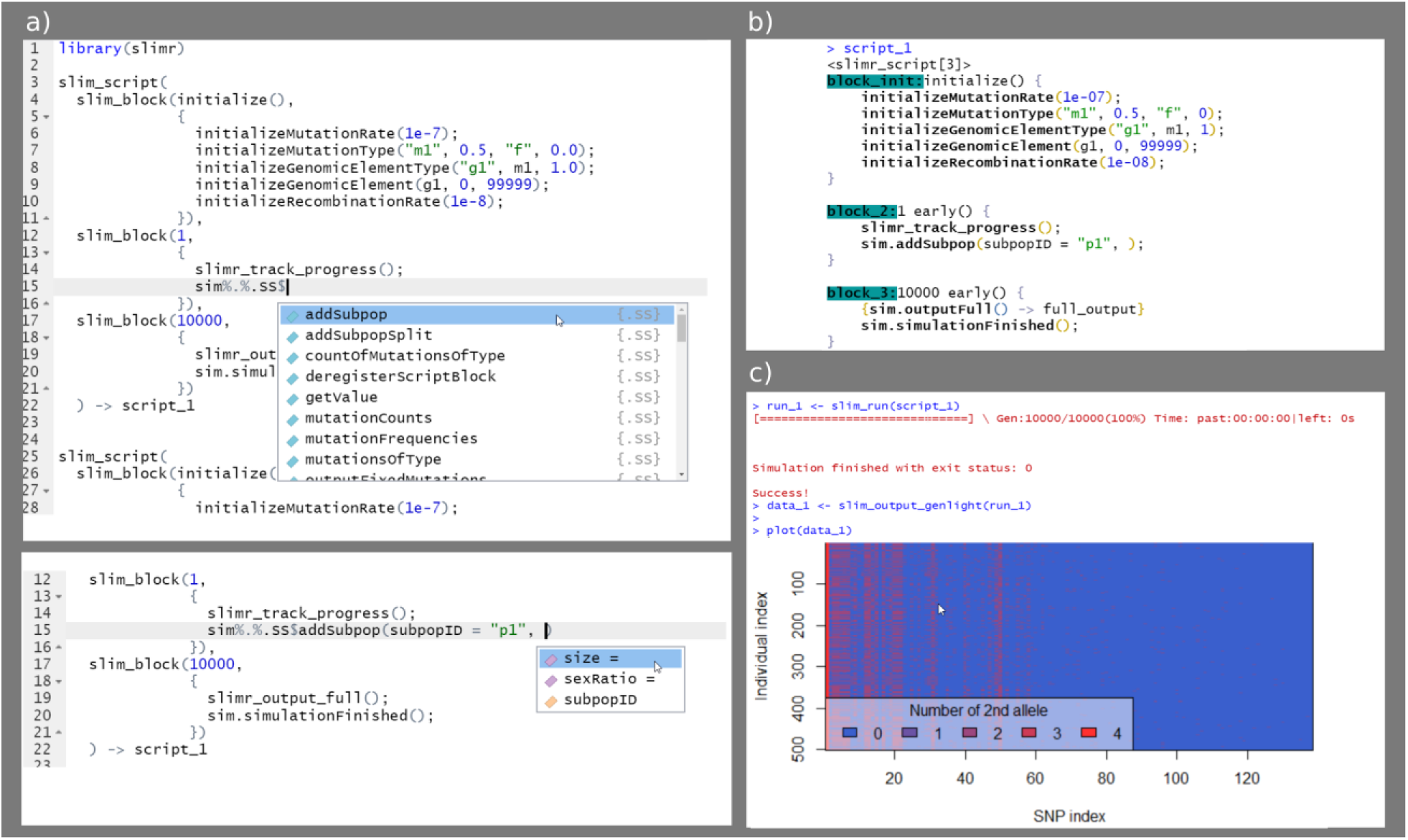
Screenshots of working with `slimr` in Rstudio. An example of autocomplete for SLiM code (a), an example of pretty printing of a `slimr_script` object (b), and an example of running a script, converting its output to a standard R format for genetic data (adegenet package’s genlight), and plotting it.

`slimr` makes it easy to write SliM code in R after the user learns a few differences between SliM and slimr. This means users can learn how to write complex SliM simulations by reading SliM 3.0 documentation and the examples found within it (https://messerlab.org/slim/). To make this process easier for `slimr` users, the entire reference documentation for functions in SliM 3.0 and Eidos scripting language (on which SliM is based) is included in `slimr` (with the original author’s permission). Hence, not only can R users look up relevant SliM functions in their R session, but the R IDE can perform autocompletion.

`slimr` also provides several Rstudio addins to make common tasks simple. These include an addin that converts SliM code to `slimr` code automatically by pasting from the clipboard, and an addin to easily send `slimr_script` calls to be run in SliM. Converting code from `slimr` to SliM is also supported, including the ability to open the converted code in SLiMGUI or QtSLiM if installed.

`slimr_script` objects (and `slimr_script_coll`, which contain lists of `slimr_script` objects) can be run in SliM, and their results collected and returned using the `slim_run` function.

### Data Input/Output

The input/output and metaprogramming features of `slimr` are achieved using special `slimr` “verbs” that can be inserted directly into `slimr` coding blocks (Fig. 2). These verbs are pure R functions that modify how the SliM script will be generated and run in SliM. They are not passed directly to SliM, but make it easy for R to interact with SliM. In this way, `slimr` code appears to be a hybrid between SliM and R code. `slimr` verbs allow all setup and logic required to use SliM with R to occur inside the coding blocks comprising the slimr_script object, thus requiring fewer arguments to be set in preparation for downstream analysis (for example `slim_run` does not require many complex arguments because most of what it needs to know is embedded in the `slimr_script` object). In our experience, this leads to a very smooth experience using `slimr` by reducing the frequency of switches between different mental modes. By convention, all `slimr` verbs have the prefix `slimr_`, and are meant to be used only inside `slim_script` calls. All other `slimr` functions are prefixed with `slim_`, which means they are to be used on `slimr_script` objects, and not inside a `slim_script` call. The following are the main `slimr` verbs supported:

**Figure 2.**
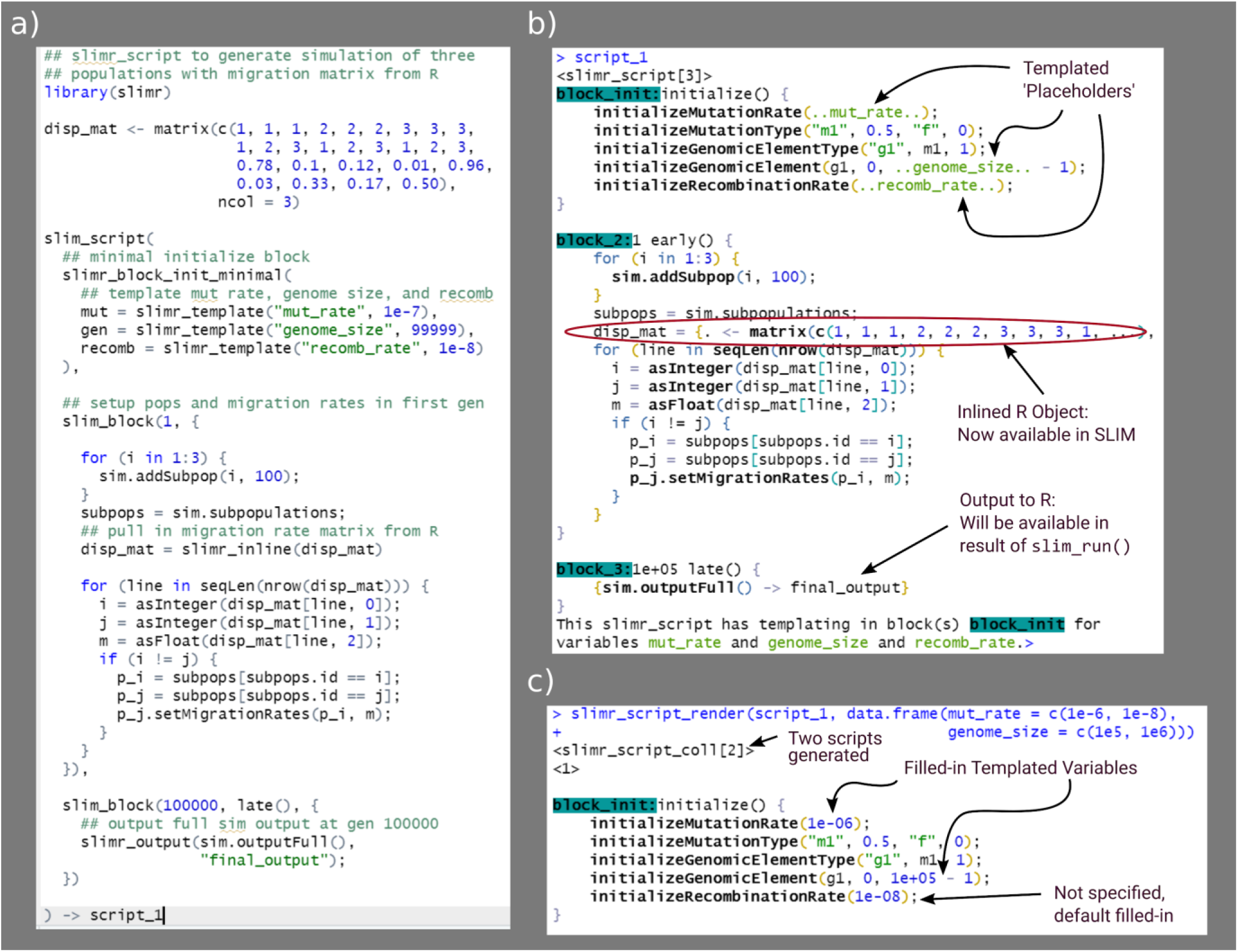
Example of a single script using the main `slimr` verbs (`slimr_template`, `slimr_output`, and `slimr_inline`). A) Code to specify the `slimr_script`. B) Pretty printing of the script, showing special `slimr` syntax. C) Example of running `slim_script_render` on the `slimr_script` object, demonstrating how placeholder variables specified in slimr_template are replaced with provided values. All code from the above example can be accessed as a package vignette (https://rdinnager.github.io/slimr/articles/simple_example.html).

#### slimr_inline

`slimr_inline` allows slimr users to embed (or “inline”) an R object directly into a SLiM script so that it can be accessed from within a SliM simulation. This is a powerful way to use empirical data that has been generated, loaded, and / or cleaned from within R within a simulation. `slimr_inline` automatically detects the type of R object and attempts to coerce it into a format compatible with SLiM. Currently supported types are all atomic vectors, matrices, arrays, and Raster* objects from the raster package, which will allow users to insert maps for use in spatial simulations.

#### slimr_output

`slimr_output` makes it simple to output data from the simulation by wrapping a SLiM expression. Where it is called in the `slimr_script,` it will produce SLiM code to take the output of the expression and send it to R. The output will be available after running `slim_run` in the returned object as a data.frame. Output can even be accessed live during the simulation run via the use of callback functions. A do_every argument tells `slimr_output` not to output every time it is called, but rather only after every `do_every` generations.

`slimr` includes several functions to create different commonly desired outputs and visualizations, which use the `slimr_output_` prefix (e.g. `slimr_output_nucleotides(`), which outputs DNA sequences data for nucleotide-based simulations).

### Metaprogramming

Metaprogramming is programming that generates or manipulates programming code itself. `slimr` has facilities for manipulating SLiM programming code and generating scripts. The main `slimr` verb for doing this is `slimr_template`. `slimr` also supports the metaprogramming operators for forcing (!!) and splicing (!!!), as used in the `{rlang}` R package. Here we briefly describe `slimr_template`, designed to help users easily generate many versions of a slimr_script with different parameters.

#### slimr_template

`slimr_template` allows the user to insert “templated” variables into a `slimr_script;` the call to `slimr_template` will be replaced in the SliM script with a placeholder `var_name` chosen by the user. This placeholder can be replaced with values of the user’s choice by calling `slim_script_render(slm_scrpt, template = tmplt)`, and providing a template – a list or data.frame containing values with names matching `var_name`. This action can be performed on multiple `slimr_template` variables simultaneously, as well as producing multiple replicate scripts with different combinations of replacements. This feature can create a swathe of parameter values to be run (automatically) in parallel to explore parameter space, conduct sensitivity analyses, or fit data to simulation output using methods requiring many simulation runs, such as Approximate Bayesian Computation (ABC). Users can provide a default value for each templated variable, which will be used if the user does not specify a replacement for that variable.

These features together make `slimr` far more than a simple wrapper for SliM – its goal is to enhance and complement SliM by creating a hybrid domain specific language for R. We plan to continue to increase integration of our package with SliM, and to continuously update it as new SliM versions are released in the future.

#### slim_run

Once a `slimr_script` or `slimr_script_coll` object has been created, with all SLiM simulation logic and `slimr` verbs for interacting with R, it can be sent to the SLiM software to be run using the `slim_run` function. To access this functionality, users must install SLiM on their computers. This is facilitated by the `slim_setup()` function, which will attempt to automatically install a platform appropriate version of SLiM on the user’s system and set it up to work with `slimr`.

Calling `slim_run` will run the simulation. While the simulation is running, `slimr_run` produces progress updates if requested, as well as any output generated by calls to `slimr_output` with custom callbacks. If called on a `slimr_script_coll` containing multiple `slimr_script` objects, each `slimr_script` object will be run, optionally in parallel, and the result returned in a list.

Once finished, `slim_run` wil return a `slimr_results` object, which contains information about the simulation run, such as whether it succeeded or failed, any error messages produced, all output generated from slimr_output calls, and any file names where additional data from the run are stored. This can then be used for any downstream analysis the user desires.

### Examples

Here we demonstrate the use `slimr` a short and simple example, and one more extensive example.

#### Simulating Nucleotide Evolution

The following script simulates a population 100 individuals that randomly splits into two equally sized subpopulations at some rate (with a probability split_prob in each generation). It simulates genomic evolution with an explicit nucleotide sequence evolution model (Jukes-Cantor model). By default SLiM only simulates and keeps track of ‘mutations’ in a more abstract sense (these could be thought of as generating new alleles at a gene, or SNPs, or however the researcher wants to interpret them). This example demonstrates the easiest way to get data from R into a slimr simulation, by using the forcing operator !!. The forcing operator tells R to evaluate what comes after first, and insert the result in its place (hence the term forcing: it forces early evaluation, normally R doesn’t evaluate variables until they are used). In the script below, we have highlighted where this is occurring in bold.

~~~
## set some parameters
seed <- 1205
split_prob <- 0.001
max_subpops <- 10
## specify simulation
split_isolate_sim <- slim_script(
slim_block(initialize(), {
  setSeed(**!!seed**);
  ## tell SLiM to simulate nucleotides
  initializeSLiMOptions(nucleotideBased=T);
  initializeAncestralNucleotides(randomNucleotides(1000));
  initializeMutationTypeNuc(“m1”, 0.5, “f”, 0.0);
  initializeGenomicElementType(“g1”, m1, 1.0, mmJukesCantor(1e-5));
  initializeGenomicElement(g1, 0, 1000 - 1);
  initializeRecombinationRate(1e-8);
}),
slim_block(1, {
  defineGlobal(“curr_subpop”, 1);
  sim.addSubpop(curr_subpop, 100)
}),
slim_block(1, 10000, late(), {
  if(rbinom(1, 1, **!!split_prob**) == 1) {
    ## split a subpop
    subpop_choose = sample(sim.subpopulations, 1)
    curr_subpop = curr_subpop + 1
    sim.addSubpopSplit(subpopID = curr_subpop,
                       size = 100,
                       sourceSubpop = subpop_choose)
    ## if too many subpops, remove one randomly
    if(size(sim.subpopulations) > **!!max_subpops**) {
      subpop_del = sample(sim.subpopulations, 1)
      subpop_del.setSubpopulationSize(0)
    }
  }
  ## output nucleotide data
  slimr_output_nucleotides(subpops = TRUE, do_every = 100)
}),
slim_block(10000, late(), {
  sim.simulationFinished()
})
)
results <- slim_run(split_isolate_sim)
~~~

The data looks like this:

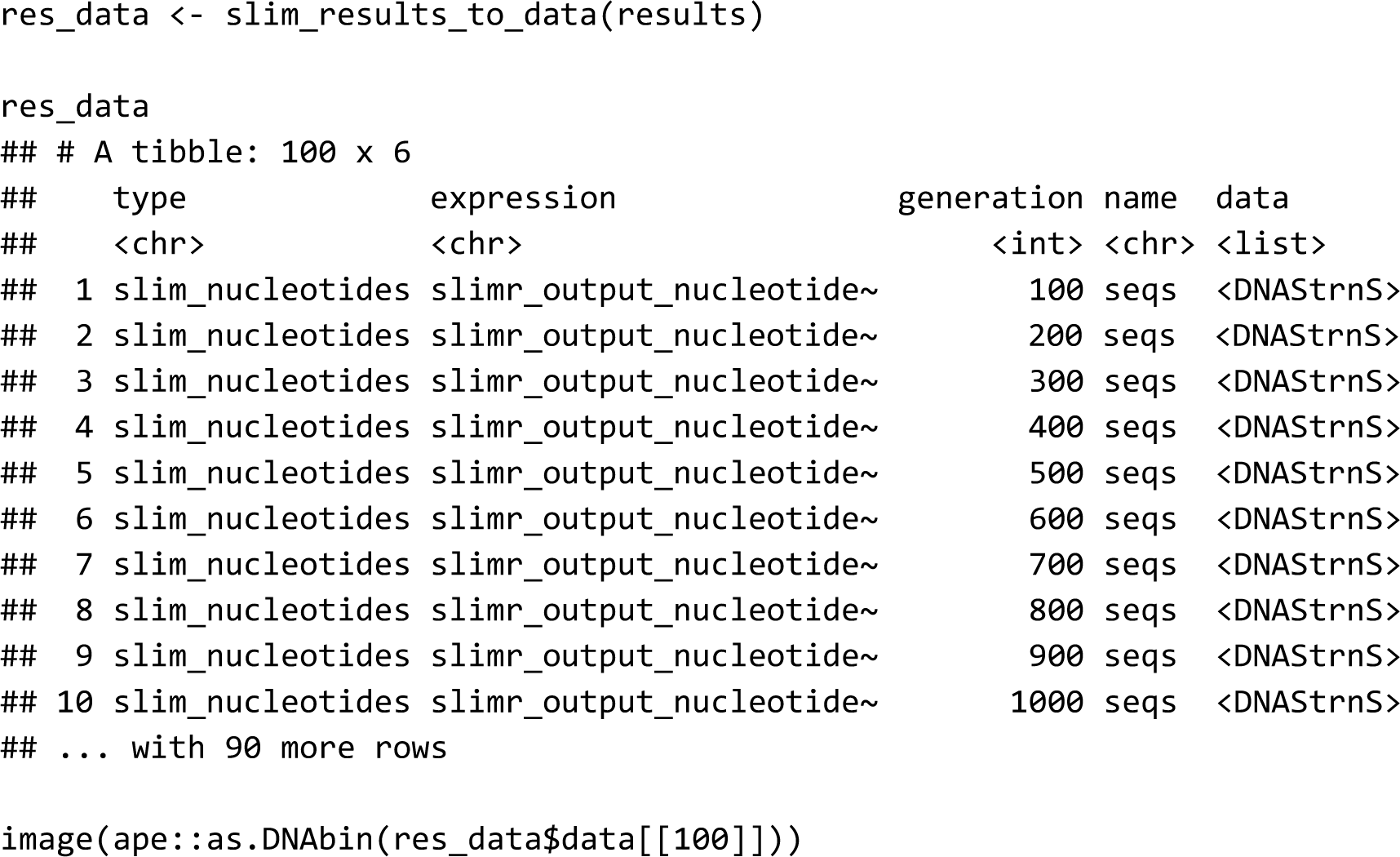

And then we can use some other R packages to quickly build a tree based on the simulated nucleotides, to see if it looks like what we would expect from a sequentially splitting population.

~~~
## convert to ape::DNAbin
al <- ape::as.DNAbin(res_data$data[[100]])
dists <- ape::dist.dna(al)
upgma_tree <- ape::as.phylo(hclust(dists, method = “average”))
pal <- paletteer::paletteer_d(“RColorBrewer::Paired”, 10)
plot(upgma_tree, show.tip.label = FALSE)
ape::tiplabels(pch = 19, col = pal[as.numeric(as.factor(res_data$subpops[[100]]))])
~~~

### Scientific Hypothesis Exploration Example: Investigating population genomics of small mammals in a periodic environment

In this section we provide a brief description of a full example analysis using simulation (Fig. 3, Fig. 4). Full code for the example can be found in the supplementary material.

**Figure 3.**
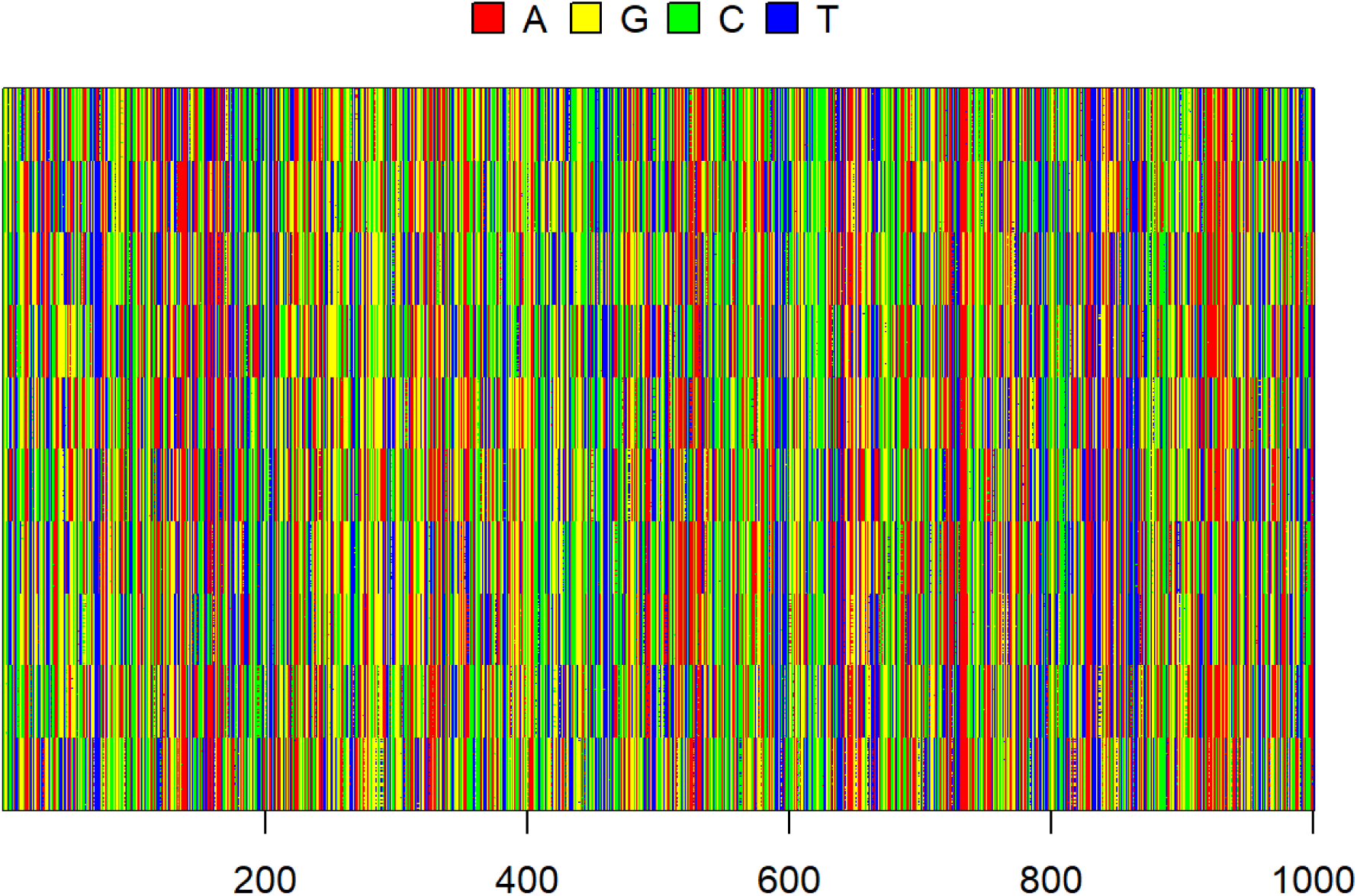
Simulated sequences for each individual. Subpopulation clustering is obvious.

**Figure 4.**
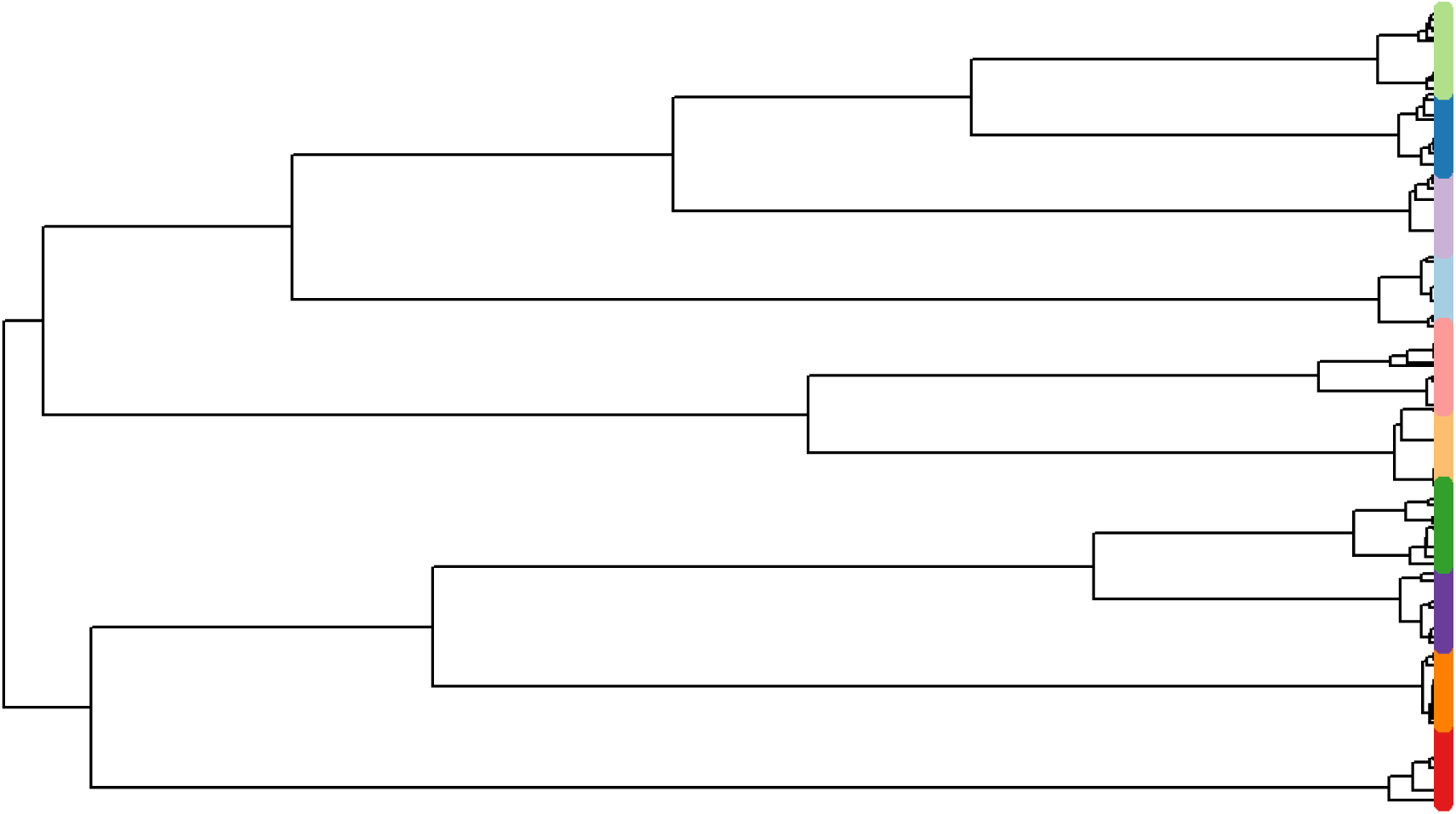
UPGMA tree of simulated subpopulations, tip points coloured by subpopulation.

The context for this example is a long-term ecological study in the Simpson Desert in central Australia. Several authors of this paper have studied the population dynamics of small mammals and reptiles in this desert for more than 30 years (C. Dickman, Wardle, Foulkes, & de Preu, 2014; Greenville, Dickman, & Wardle, 2017; Greenville, Wardle, Nguyen, & Dickman, 2016). Recently, we have begun sequencing tissue samples taken from animals captured during the past 15 years, and obtained single nucleotide polymorphism (SNP) data using DArT (Diversity Arrays Technology Pty Ltd) technology. Here, we use SNP data from 167 individuals of a common native rodent species, the sandy inland mouse *Pseudomys hermannsburgensis*, sampled at 7 sites over three years (2006-2008), and subsequently aggregated to 3 subpopulations for analysis. The three sample years span periods before and after a major rainfall event at the end of 2006; big rains occur infrequently in the study region (every 8-12 years) (Greenville, Wardle, & Dickman, 2012) but drive major population eruptions.

We used the SNP data to calculate pairwise F_st_ values among the three subpopulations in each year, revealing that pairwise F_st_ values dropped rapidly to nearly zero immediately after the rainfall event from a high recorded just prior to the event when the populations were more genetically differentiated. We interpreted this result to mean that the rainfall event, which caused the sandy inland mouse population to rapidly increase, also allowed animals to move out of spatially scattered refuge patches to which they had been confined during the preceding dry period (C. R. Dickman, Greenville, Tamayo, & Wardle, 2011). This movement allowed the subpopulations to mix, leading to a decrease in population genetic structure as measured by F_st_.

In the example, we use simulations to evaluate our interpretation regarding the processes driving changes in F_st_ values (Figure 5). We found that our initial hypothesis, that the rainfall event led to the mixing of previously unconnected populations in refuge patches, provided a good match to the data when we simulated the population and genomic processes (Figure 6). However, we also identified several other processes that could generate similar outcomes, which raises the question as to what data or analyses would be required to distinguish among these competing processes.

**Figure 5.**
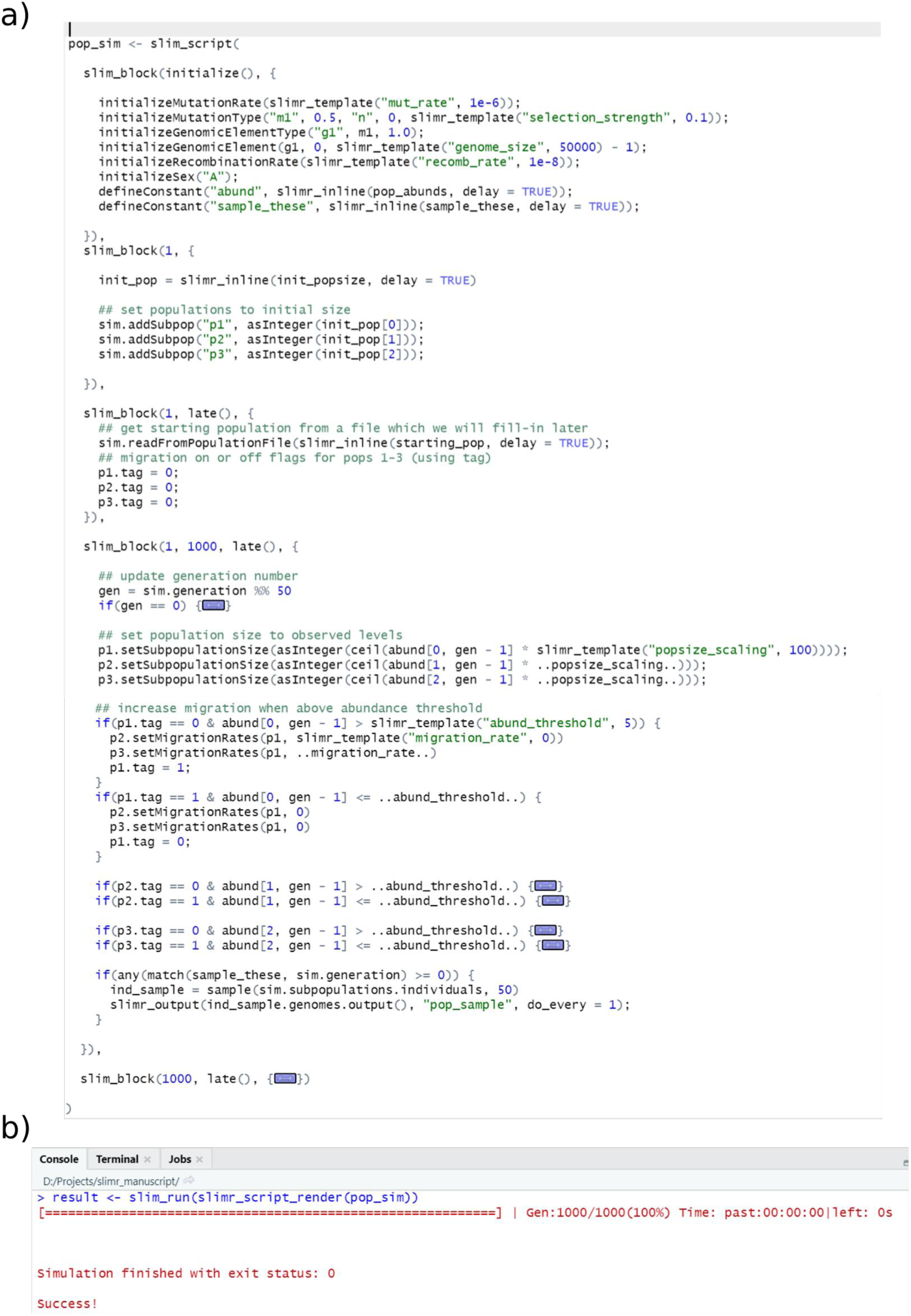
Screenshots of the main `slimr` example from this manuscript. **a)** Rstudio screenshot showing completed slimr script for specifying the model. **Note:** some code sections have been collapsed for brevity. Full code can be found in the Supplementary Material. **b)** Screen shot of the Rstudio console after running the example using default values for templated variables. This shows how fast SLiM can run – this example took less than 1 second!

**Figure 6.**
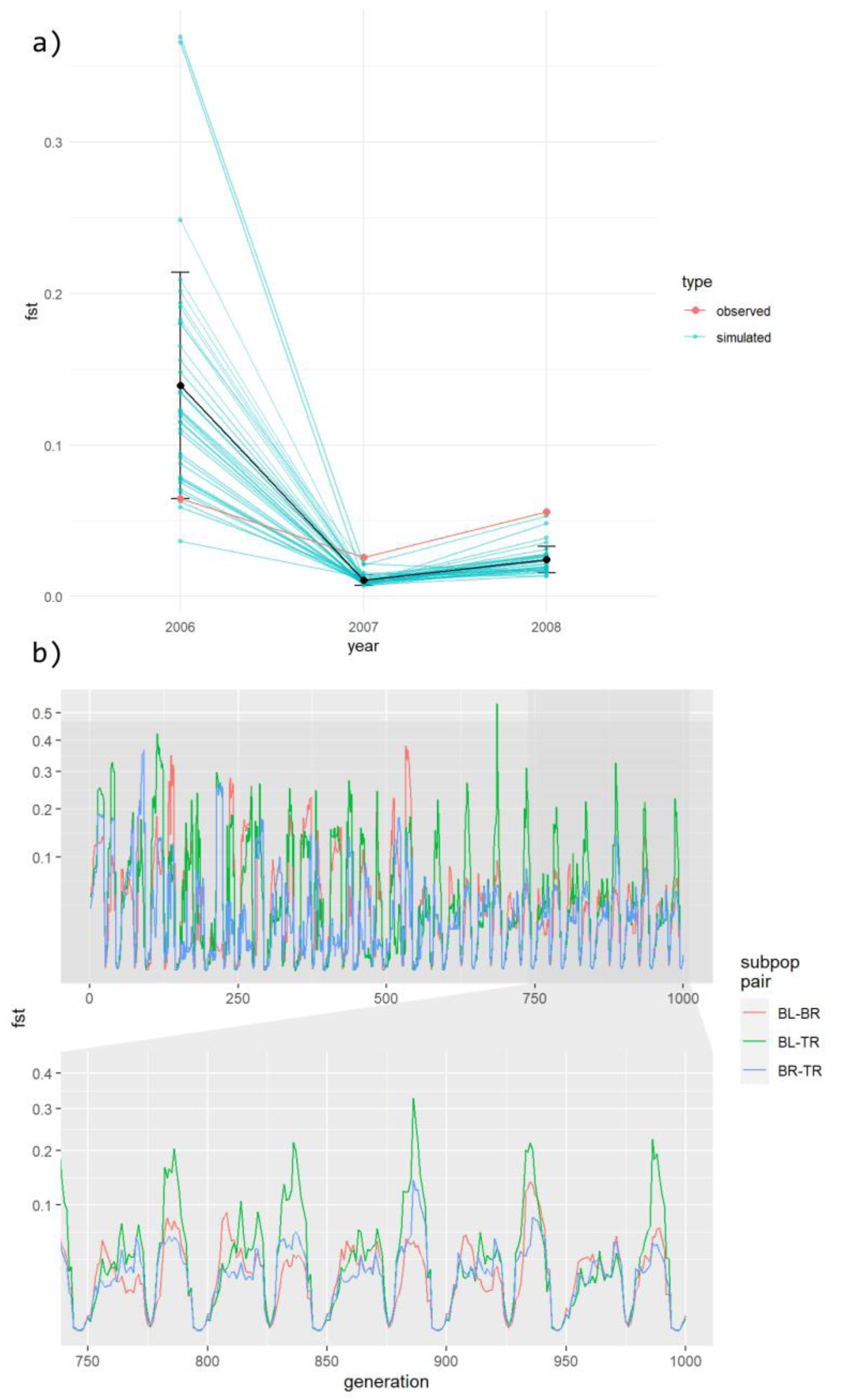
**A)** Mean F_st_ values from 36 replicate simulations simulated under our hypothesized mechanism to explain Fst fluctuations, using hand chosen parameter values. Blue values and lines represent simulated values, red values and line represents the observed F_st_ values. Details of simulation including code is in the Supplementary material. **B)** Same simulation run over many generations, showing the three subpopulations separately.

To formalize our ideas a little more we ran an Approximate Bayesian Computation (ABC) analysis to derive an approximate posterior distribution of model parameters that produced a good fit to our short F_st_ time series (see Supplementary Materials: ABC Analysis for code used). We were able to easily move from simulation exploration to a more formal fitting exercise because the simulation was already in R (thanks to `slimr`), and so only a small amount of code was required to convert the input and output of our simulation to the format required by the `easyABC` package, which we used for this analysis.

### ABC Results

After extracting the parameters of a sample of the approximate posterior distribution we reran simulations based on those parameters, calculated mean Fst and plotted them next to the observed Fst values (Figure 7). The simulated Fsts do cluster around the observed values though it does appear that the simulated values for 2018 (the year after the rainfall) do tend to be a bit lower than the observed values in the simulations.

**Figure 7.**
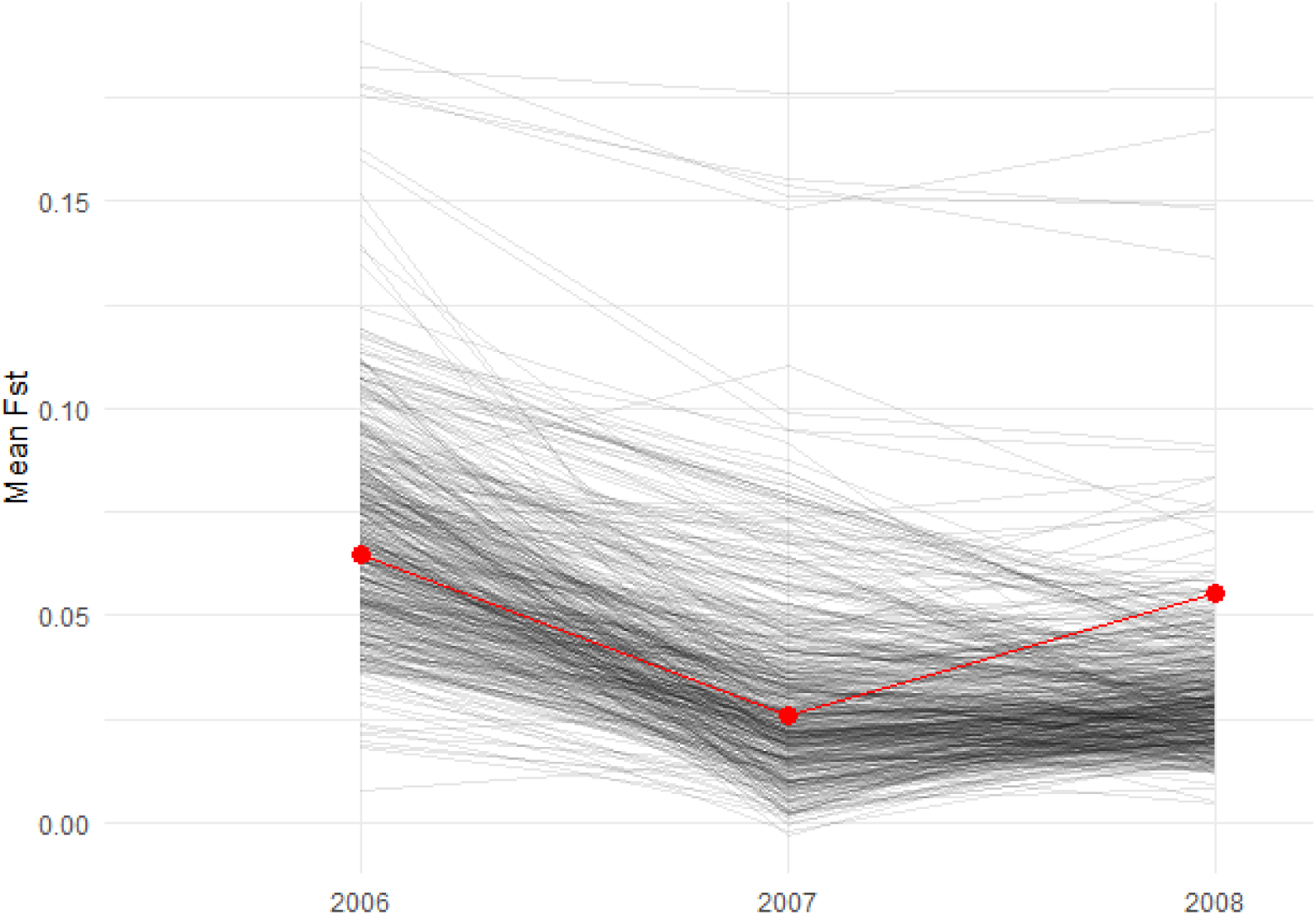
Fst values calculated from simulations based on 500 parameter value sets drawn from the approximate posterior distribution of our simulation, based on an ABC analysis. Partially transparent black line represent the simulations. Red points and line represent the observed Fst values from the study described in this section.

The marginal posterior distribution based on samples from the ABC analysis confirms that our data do not constrain individual parameters much, with a fairly wide distribution for most parameters providing a good fit to our data (Figure 8, diagonal panels). The only exception was perhaps mutation rate, for which the lower values that we simulated tended to provide a better fit. The parameter of most interest to us was the abundance threshold (abund_threshold in Figure 8), which specified the population size above which a subpopulation would ‘turn on’ migration, that is, start exporting individuals to the other subpopulations (in the real system this population size change is driven by rainfall). In this simulation an abundance threshold of zero or less would be migration always happening, and one of 20 or over would be migration almost never happening. Some simulations produced well fitting Fst values for nearly all relevant values of the abundance threshold, with some falloff at either end. However, when we start looking at combinations of multiple parameters we see that the value of the abundance threshold parameter does constrain what values of other parameters will make for a good fit to the data. For example, if the abundance threshold is low, and thus migration is always on, only simulations with very low migration rates and very low mutation rates can provide a good fit to the data (figure 8, panels in rows 3 and 4 in column 2). All in all this suggests that there are two approaches to improving our ability to distinguish how different processes lead to the patterns we see (besides just collecting more data): 1) try adding new summary statistics besides just pairwise Fst, which may capture some other aspect of the data, and 2) use some independent sources of data or information to estimate and constrain the parameter space of our simulations closer to that of the real system. In particular, approach 1 could be tested without having to collect more data by doing more simulations: we could simulate our model, then simulate data collection and calculate our new summary statistic on the simulated data. We can then see if we can recover the parameters of our simulation better than we could before after incorporating our new statistic.

**Figure 8.**
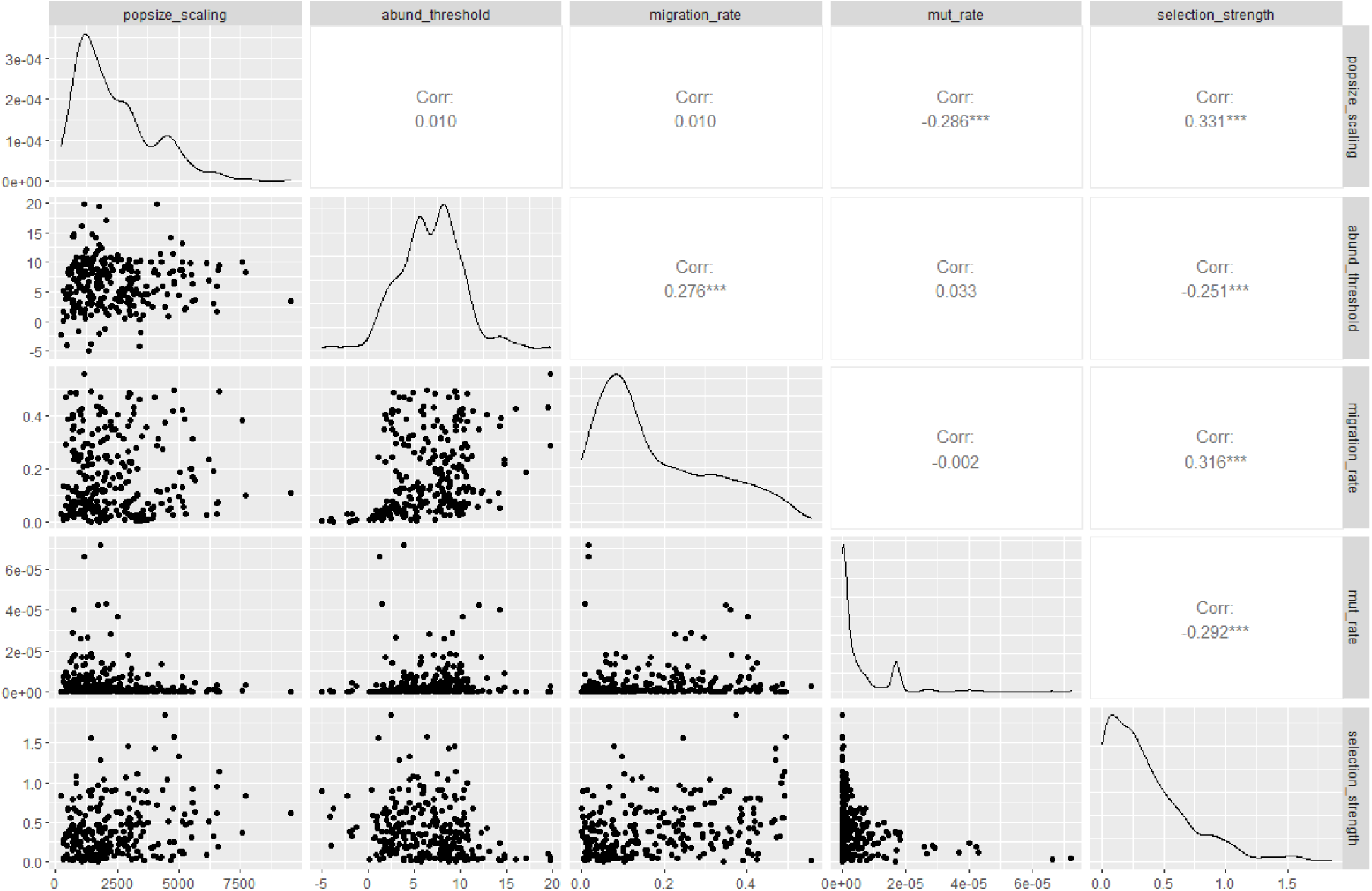
Samples from the approximate posterior distribution for our simulation model fit with Approximate Bayesian Computation. Lower left panels show the 5 parameters of our simulation model estimated, plotted against each other in a pairwise fashion, giving an indication of their joint posterior distribution. Some parameters are highly correlated in the posterior. Upper right panels show the pearson correlation coefficient, and the panels in the diagonal show the marginal distribution of each parameter estimated using kernel density estimation on the samples.

The results from these preliminary simulations will thus be invaluable in guiding which individuals and time periods we should focus our sequencing on, and what summary statistics to use, to maximize the chances of distinguishing among competing hypotheses that might explain the combined population and genetic patterns in the data. Ultimately, we aim to use this approach to understand how future climate change could alter the population and genetic structure of desert animals, highlighting the value of `slimr` in a scientific workflow.

### slimr and Open Science

It is increasingly being seen as vital for biologists to share code used to generate their results in the spirit of open science. A researcher may spend months perfecting a SLiM script that simulates a particular scenario of interest, but this scenario and those similar to it are likely of interest to other researchers as well. `slimr` allows the sharing of simulations in a very open and easy to use way, through the R software ecosystem. It provides tools that can allow researchers, with very little additional code, to make their simulations accept user-defined input, and output to common formats used by R users. Simulations can easily be wrapped into R package, which can then be installed by any R user with a command. Because `slimr` provides general interfacing functionality from SLiM to R, it allows open development of simulations by developers with much less experience with R coding, and requiring far less time.

To demonstrate this functionality, and to provide an alternative way for simulation developers to share their simulations if they do not have the time or experience to write an entire package, we have developed a companion R package called `slimrmodels` (https://github.com/rdinnager/slimrmodels), in which we have implemented several potentially useful simulation models as user-friendly functions, including the simulation developed for our main example in this manuscript. We freely encourage other researchers to contribute their own models to this package, by making a pull request on github. Using the functions in `slimrmodels` requires no knowledge of SLiM code to use, and completely hides SLiM from users.

Nevertheless, all models can be exported as a SLiM script for further customizations by users knowledgeable in SLiM. `slimrmodels` is released under an MIT license, and will be continuously contributed to by our research group and (we hope) other research groups, as new models are developed.

In `slimrmodels` a run of the main example simulation used here (Figure 5) can be coded succinctly as:

~~~
results <- slimrmodels::mod_fixed_pop_dyn(
             pop_abund = function(gen, pop_scale, …) pop_values * pop_scale,
             sampler = samp_these,
             migration_rates = function(gen, abund_threshold, mig_rate, …) {
                    ifelse(pop_values > abund_threshold, mig_rat, 0)
             },
             pop_scale = pop_scale,
             abund_thres = abund_thres,
             mig_rate = mig_rate)
~~~

More information about `slimrmodels` can be found in the documentation for the package (https://github.com/rdinnager/slimrmodels).

## Supporting information

Main example vignette and code

## Acknowledgments

We thank Benjamin C. Haller and Phillip W. Messer for permission to reproduce the documentation and examples of SLiM in `slimr`, and for valuable feedback on the package (from B.C.H.). Thanks to Emily Stringer for providing additional test data to help develop the methods used in this manuscript.

## Author Contributions

RD, BG, SS, and RPD developed the concept for the package. SE, CD, GW, and AG provided feedback on the package design. RD coded the package and wrote the manuscript draft. CD, GW, and AG contributed data for testing of the package, and BG helped test the package as a user. All authors contributed critically to manuscript drafts and gave final approval for publication.

