## Supplementary material for "slimr: An R package for integrating data and tailor-made population genomic simulations over space and time": Main example vignette and code: ABC_example.R

```
## This script sets up a simple 3 subpopulation simulation and then 
## fits its parameters using ABC

library(slimr)
library(EasyABC)
library(readr)

## load starting data
pop_abunds <- read_rds("data/pop_abunds.rds")
samp_sizes <- read_rds("data/samp_sizes.rds")
init_popsize <- read_rds("data/init_popsize.rds")

starting_pop = "sims/starting_pop.txt" %>%
  slim_file()

## load target Fst values
fst_df <- read_rds("data/fst_df.rds")
target_fsts <- fst_df$fst

default_genome_size <- 300000
## set sample times corresponding to our genetic data (roughly)
sample_times <- c("2006" = 40, "2007" = 45, "2008" = 49)
sample_these <- purrr::map(c(4:9),
                           ~50*.x + sample_times) %>%
  purrr::reduce(union)

## setup functions:
sim.mutationFrequencies <- function(subpop) {
  gen <- get("gen", parent.frame(2))
  gl.alf(gen[pop = subpop])[ , 2]
}

isFinite <- function(x) is.finite(x)

calculateFST <- function(inds1, inds2) {
  ## Calculate the FST between two subpopulations
  p1_p = inds1.genomes.mutationFrequenciesInGenomes();
  p2_p = inds2.genomes.mutationFrequenciesInGenomes();
  mean_p = (p1_p + p2_p) / 2.0;
  H_t = 2.0 * mean_p * (1.0 - mean_p);
  H_s = p1_p * (1.0 - p1_p) + p2_p * (1.0 - p2_p);
  fst = 1.0 - H_s/H_t;
  fst = fst[isFinite(fst)]; ## exclude muts where mean_p is 0.0 or 1.0
  return(mean(fst));
}

pairFST <- function(gen) {
  
  pops <- as.character(unique(pop(gen)))
  pop_pairs <- combn(pops, 2)
  pair_fst <- apply(pop_pairs, 2, function(x) calculateFST(x[1], x[2]))
  fst_mat <- matrix(nrow = length(pops), ncol = length(pops))
  rownames(fst_mat) <- colnames(fst_mat) <- pops
  fst_mat[t(pop_pairs)] <- pair_fst
  t(fst_mat)
  
}

## specify simulation

slim_script(
  
  slim_function("o<Individual> inds1", "o<Individual> inds2",
                name = "calculateFST",
                return_type = "f$", body = calculateFST),
  
  slim_block(initialize(), {
    
    setSeed(slimr_template("seed", 123456))
    #initializeSLiMOptions(keepPedigrees=T);
    initializeMutationRate(slimr_template("mut_rate", 1e-6));
    initializeMutationType("m1", 0.5, "n", 0, slimr_template("selection_strength", 0.1));
    initializeGenomicElementType("g1", m1, 1.0);
    initializeGenomicElement(g1, 0, slimr_template("genome_size", !!default_genome_size) - 1);
    initializeRecombinationRate(slimr_template("recomb_rate", 1e-8));
    initializeSex("A");
    defineConstant("abund", slimr_inline(pop_abunds, delay = FALSE));
    defineConstant("sample_these", slimr_inline(sample_these, delay = FALSE));
    defineConstant("samp_sizes", slimr_inline(samp_sizes, delay = FALSE));
    
  }),
  slim_block(1, {
    
    init_pop = slimr_inline(init_popsize, delay = FALSE)
    
    ## set populations to initial size
    sim.addSubpop("p1", asInteger(init_pop[0]));
    sim.addSubpop("p2", asInteger(init_pop[1]));
    sim.addSubpop("p3", asInteger(init_pop[2]));
    
  }),
  
  slim_block(1, late(), {
    ## get starting population from a file which we will fill-in later
    sim.readFromPopulationFile(slimr_inline(starting_pop, delay = FALSE));
    ## migration on or off flags for pops 1-3 (using tag)
    p1.tag = 0;
    p2.tag = 0;
    p3.tag = 0;
  }),
  
  slim_block(1, 500, late(), {
    
    ## update generation number
    gen = sim.generation %% 50
    if(gen == 0) {
      gen = 50
    }
    
    ## set population size to observed levels
    p1.setSubpopulationSize(asInteger(ceil(abund[0, gen - 1] * slimr_template("popsize_scaling", 100))));
    p2.setSubpopulationSize(asInteger(ceil(abund[1, gen - 1] * ..popsize_scaling..)));
    p3.setSubpopulationSize(asInteger(ceil(abund[2, gen - 1] * ..popsize_scaling..)));
    
    ## increase migration when above abundance threshold
    if(p1.tag == 0 & abund[0, gen - 1] > slimr_template("abund_threshold", 5)) {
      p2.setMigrationRates(p1, slimr_template("migration_rate", 0))
      p3.setMigrationRates(p1, ..migration_rate..)
      p1.tag = 1;
    } 
    if(p1.tag == 1 & abund[0, gen - 1] <= ..abund_threshold..) {
      p2.setMigrationRates(p1, 0)
      p3.setMigrationRates(p1, 0)
      p1.tag = 0;
    }
    
    if(p2.tag == 0 & abund[1, gen - 1] > ..abund_threshold..) {
      p1.setMigrationRates(p2, ..migration_rate..)
      p3.setMigrationRates(p2, ..migration_rate..)
      p2.tag = 1;
    } 
    if(p2.tag == 1 & abund[1, gen - 1] <= ..abund_threshold..) {
      p1.setMigrationRates(p2, 0)
      p3.setMigrationRates(p2, 0)
      p2.tag = 0;
    }    
    
    if(p3.tag == 0 & abund[2, gen - 1] > ..abund_threshold..) {
      p1.setMigrationRates(p3, ..migration_rate..)
      p2.setMigrationRates(p3, ..migration_rate..)
      p3.tag = 1;
    } 
    if(p3.tag == 1 & abund[2, gen - 1] <= ..abund_threshold..) {
      p1.setMigrationRates(p3, 0)
      p2.setMigrationRates(p3, 0)
      p3.tag = 0;
    }
    
    ## only run if the generation is in our sample_these list
    if(any(match(sample_these, sim.generation) >= 0)) {
      ## find the sample size that matches the matching "year" for our obs data
      ssizes = drop(samp_sizes[ , which(sample_these == sim.generation)]) 
      ## sample individuals
      p1_samp = sample(p1.individuals, ssizes[0])
      p2_samp = sample(p2.individuals, ssizes[1])
      p3_samp = sample(p3.individuals, ssizes[2])
      p1_p2_fst = calculateFST(p1_samp, p2_samp)
      p1_p3_fst = calculateFST(p1_samp, p3_samp)
      p2_p3_fst = calculateFST(p2_samp, p3_samp)
      ## output individuals genomes
      slimr_output(p1_p2_fst, "p1_p2_fst", do_every = 1)
      slimr_output(p1_p3_fst, "p1_p3_fst", do_every = 1);
      slimr_output(p2_p3_fst, "p2_p3_fst", do_every = 1);
    }
    
  }),
  
  slim_block(500, late(), {
    sim.simulationFinished()
  })
  
) -> pop_sim_samp

pop_sim_samp

## do a test run with default parameters
test_run <- slim_run(pop_sim_samp)

dat <- slim_results_to_data(test_run)
dat
as.numeric(unlist(dat$data))

## take the mean over our 9 desired values
apply(matrix(as.numeric(unlist(dat$data)), nrow = 9), 1, mean)

## remind ourselves what the parameters are:
slimr_template_info(pop_sim_samp)

## some of these are 'nuisance parameter' and are unlikely to be identifiable,
## so we will just set them to a constant (their default value):
## recomb_rate, genome_size, the other 5 parameters we will vary

## In order to run this model with easyABC we just need to create a wrapper function
## that contains everything the easyABC functions need:

make_easyABC_fun <- function(sim) {
  function(x) {
    ## protect against numerical undeflow
    x[5] <- exp(x[5])
    suppressWarnings(
      scrpt <- slimr::slim_script_render(sim, template = list(popsize_scaling = x[2],
                                                              abund_threshold = x[3],
                                                              migration_rate = x[4],
                                                              mut_rate = x[5],
                                                              selection_strength = x[6],
                                                              seed = x[1]))
    )
    
    run <- try(suppressMessages(slimr::slim_run(scrpt, progress = FALSE, throw_error = TRUE)),
               silent = TRUE)
    
    ## this bit works around parameter values that cause the sim to fail
    ## if the simulation throws an error, return a vector of 2s (an impossible value for Fst
    ## which should be far away from the target and so should get thrown out by the ABC filter)
    if(inherits(run, "try-error")) {
      res <- rep(2, 9)  
    } else {
      dat <- slimr::slim_results_to_data(run)
      res <- apply(matrix(as.numeric(unlist(dat$data)), nrow = 9), 1, mean)
    }
    
    res
    
  }
}

run_sim <- make_easyABC_fun(pop_sim_samp)

run_sim(c(123, 100, 5, 0.1, log(1e-06), 0.1))
run_sim(c(123, 10, 5, 0.1, log(1e-06), 0.1))

## run ABC! Start with a simple sequential method

ABC_res <- ABC_sequential(method = "Lenormand", model = run_sim,
                          prior = list(popsize_scaling = c("unif", 100, 1000),
                                       abund_threshold = c("unif", 0, 20),
                                       migration_rate = c("unif", 0, 0.5),
                                       mut_rate = c("unif", log(1e-07), log(1e-05)),
                                       selection_strength = c("unif", 0, 0.5)),
                          summary_stat_target = target_fsts,
                          nb_simul = 500,
                          use_seed = TRUE,
                          progress_bar = interactive(),
                          n_cluster = 20,
                          verbose = TRUE,
                          p_acc_min = 0.05,
                          inside_prior = TRUE)

write_rds(ABC_res, "data/ABC_res.rds")

ABC_res_big <- ABC_sequential(method = "Lenormand", model = run_sim,
                          prior = list(popsize_scaling = c("unif", 100, 2000),
                                       abund_threshold = c("unif", 0, 20),
                                       migration_rate = c("unif", 0, 0.5),
                                       mut_rate = c("unif", log(1e-07), log(1e-05)),
                                       selection_strength = c("unif", 0, 0.5)),
                          summary_stat_target = target_fsts,
                          nb_simul = 1000,
                          use_seed = TRUE,
                          progress_bar = interactive(),
                          n_cluster = 20,
                          verbose = TRUE,
                          p_acc_min = 0.01,
                          inside_prior = FALSE)

write_rds(ABC_res_big, "data/ABC_res_big.rds")
```
