## Supplementary material for "slimr: An R package for integrating data and tailor-made population genomic simulations over space and time": Main example vignette and code: Main_manuscript_example_v2.html

Using slimr to investigate the population genetics of the Sandy Inland Mouse in the periodic rainfall environment of the Simpson Desert


### Using slimr to investigate the population genetics of the Sandy Inland Mouse in the periodic rainfall environment of the Simpson Desert

###### Russell Dinnage

#### 09/02/2022

#### A look at the data

An ongoing project in the Simpson Desert of central Australia is producing genomic sequence data for small mammals and reptiles collected over more than 30 years. We have some preliminary data from this project for the Sandy Inland Mouse (*Pseudomys hermannsburgensis*) that we’d like to use to explore, through data analysis and simulation, and hopefully gain some insight into where this project might ultimately take us. First let’s load all the packages we use in the example, and then load the genetic data we will be working with. The data is stored in an `.Rdata` file containing a `genlight` object, which stores binary Single Nucleotide Polymorphism (SNP) data, defined in the `adegenet` package. We also have demographic data in the form of trap capture data (captures per 100 trap nights; see Dickman et al. 2014 and Greenville et al. 2016 for capture methods).

```
library(readr)
library(dplyr)
library(tidyr)
library(tibble)
library(mapview)
library(purrr)
library(conflicted)
library(future)
library(furrr)
library(lubridate)
library(ggplot2)
library(ggforce)
library(gganimate)
library(directlabels)
library(patchwork)
library(sf)
library(adegenet)
library(dartR)
library(here)
library(stringr)
library(slimr)

mapviewOptions(fgb = FALSE)

## because filter function exists in multiple packages we use
## the conflicted package to make sure we R uses the version in dplyr
conflict_prefer("filter", "dplyr") 
conflict_prefer("select", "dplyr")
conflict_prefer("initialize", "slimr")

## load data
gen <- read_rds("data/herm.rdata")

gen
```

```
##  /// GENLIGHT OBJECT /////////
## 
##  // 167 genotypes,  39,978 binary SNPs, size: 28.4 Mb
##  1028248 (15.4 %) missing data
## 
##  // Basic content
##    @gen: list of 167 SNPbin
##    @ploidy: ploidy of each individual  (range: 2-2)
## 
##  // Optional content
##    @ind.names:  167 individual labels
##    @loc.names:  39978 locus labels
##    @loc.all:  39978 alleles
##    @position: integer storing positions of the SNPs
##    @pop: population of each individual (group size range: 2-39)
##    @other: a list containing: loc.metrics  latlong  ind.metrics  loc.metrics.flags  verbose  history
```

```
abund <- read_csv("data/mammal_captures.csv")

abund
```

So we have SNP sequences from 167 individuals taken over three years and several different sites. Before we take a look at the genetic data, let’s set the environmental context a bit. These small mammals live in the Simpson Desert, an arid ecosystem characterized by long periods of little rain, with occasional major rainfall events. These tend to occur every 6-10 years. After rainfall events, the desert turns green, it can be seen from space, and our analysis of satellite reflectance data confirms massive spikes in productivity after these rainfalls. Plants grow, flower and seed massively during these periods, and so food is abundant for small mammals. We can see the regular pulse of rainfall just by looking at the mammal abundances – *Pseudomys hermannsburgensis* in particular responds quickly to rainfall. Let’s look at our live trapping data on abundances to see this pattern.

```
## give more descriptive names to sites
sites <- tribble(~pop, ~SiteName,
                 "CS", "Carlo",
                 "FRN", "Field River North",
                 "FRS", "Field River South",
                 "KSE", "Kunnamuka Swamp East",
                 "MC", "Main Camp",
                 "SS", "South Site",
                 "WS", "Way Site"
)

abund_summ <- abund %>%
  select(-`Notomys alexis`, -`Sminthopsis youngsoni`) %>%
  left_join(sites) %>%
  drop_na(pop) %>%
  ## convert month-year character column to proper dates using lubridate
  separate(MonthYear, c("Month", "year"), "\\.") %>%
  mutate(Month = sapply(Month, function(x) which(month.abb == x))) %>%
  mutate(date = myd(paste(Month, year, sep = "-"), truncated = 1)) %>%
  ## summarise by site and date
  group_by(date, pop) %>%
  summarise(abund = mean(`Pseudomys hermannsburgensis`),
            .groups = "drop")
```

```
## Joining, by = "SiteName"
```

```
ggplot(abund_summ, aes(date, abund)) +
  geom_path(aes(colour = pop)) +
  geom_dl(aes(label = pop), method = "last.qp") +
  aes(colour = pop) +
  theme_minimal() +
  theme(legend.position = "none")
```

Wow, those pulses are quite clear and quite extreme!

Okay, now let’s go have a look at the SNP data we have for these populations.

First, we will extract the metadata for the genetic samples and have a look at how they are distributed in the Simpson Desert.

```
gen_meta <- gen@other$ind.metrics

gen_meta <- gen_meta %>%
  as_tibble() %>%
  ## fix some minor errors in genetic metadata with respect to site names:
  mutate(pop = case_when(pop == "FR" ~ substr(as.character(SiteGrid), 1, 3),
                         pop == "KS" ~ "KSE",
                         TRUE ~ as.character(pop))) %>%
  left_join(sites)
```

```
## Joining, by = "pop"
```

```
gen_meta
```

Extract and plot site coordinates:

```
coords <- gen_meta %>%
  group_by(pop) %>%
  summarise(lon = mean(lon),
            lat = mean(lat),
            .groups = "drop") %>%
  sf::st_as_sf(coords = c("lon", "lat"), crs = 4326)

mapview(coords, label = coords$pop, map.types = "Esri.WorldImagery")
```

Looks like our sites cluster into roughly 3 spatial groups. For simplicity we can merge these sites into 3 subpopulations, which will make it a bit simpler for our simulation later.

```
coords_3 <- gen_meta %>%
  mutate(three_pop = case_when(pop %in% c("MC", "SS", "WS") ~ "BR",
                               pop %in% c("FRN", "FRS") ~ "BL",
                               pop %in% c("KSE", "CS") ~ "TR")) %>%
  group_by(three_pop) %>%
  summarise(lon = mean(lon),
         lat = mean(lat),
         .groups = "drop") %>%
  ungroup() %>%
  sf::st_as_sf(coords = c("lon", "lat"), crs = 4326)

mapview(coords_3, label = coords_3$three_pop, map.types = "Esri.WorldImagery")
```

As it happens these merged subpopulations are roughly equidistant, arranged in a triangle. If we assume the terrain between them is similarly easy to traverse we can simplify our model of the system even more and assume that all else being equal these subpopulations should have roughly similar migration rates between them. We called these subpopulations “BR”, “BL”, and “TR”, for Bottom Right, Bottom Left, and Top Right (creatively named I know).

Given this setup, let us have a look at the Fst values between these three subpopulations over the three years. We will use a custom function to calculate pairwise Fst for our subpopulations. We will do it this way so we can show some `slimr` functionality later, where we can exploit the similarity of R and SLiM code to run our Fst function from within SLiM, and get fast and live Fst estimates while a simulation is running. This means that the Fst calculated for our observed and our simulated data will be directly comparable when we do our downstream analysis.

```
sim.mutationFrequencies <- function(subpop) {
  gen <- get("gen", parent.frame(2))
  gl.alf(gen[pop = subpop])[ , 2]
}

isFinite <- function(x) is.finite(x)

calculateFST <- function(subpop1, subpop2) {
                  ## Calculate the FST between two subpopulations
                  p1_p = sim.mutationFrequencies(subpop1);
                  p2_p = sim.mutationFrequencies(subpop2);
                  mean_p = (p1_p + p2_p) / 2.0;
                  H_t = 2.0 * mean_p * (1.0 - mean_p);
                  H_s = p1_p * (1.0 - p1_p) + p2_p * (1.0 - p2_p);
                  fst = 1.0 - H_s/H_t;
                  fst = fst[isFinite(fst)]; ## exclude muts where mean_p is 0.0 or 1.0
                  return(mean(fst));
}

pairFST <- function(gen) {
  
  pops <- as.character(unique(pop(gen)))
  pop_pairs <- combn(pops, 2)
  pair_fst <- apply(pop_pairs, 2, function(x) calculateFST(x[1], x[2]))
  fst_mat <- matrix(nrow = length(pops), ncol = length(pops))
  rownames(fst_mat) <- colnames(fst_mat) <- pops
  fst_mat[t(pop_pairs)] <- pair_fst
  t(fst_mat)
  
}
```

We used several custom functions above and named them so that they would match available functions in SLiM. That way, when we run our simulations using `slimr`, SLiM will use its optimized versions of these functions internally. We also create a function to run the pairwise Fst function on all possible subpopulation pairs, which we will now run on our observed SNP data. We used a function in the `dartR` package to calculate allele frequencies to use in our Fst calculation. In SLiM, this is a builtin function, which we will take advantage of later. Note that the use of `get` and `parent.frame` is a trick to get R to use the right `gen` object without having to specify it as an argument, which will let us reuse the functions later in SLiM, which cannot access R objects while it is running.

```
gen_meta <- gen_meta %>%
  mutate(three_pop = case_when(pop %in% c("MC", "SS", "WS") ~ "BR",
                               pop %in% c("FRN", "FRS") ~ "BL",
                               pop %in% c("KSE", "CS") ~ "TR"))

gen@other$ind.metrics <- gen_meta 
pop(gen) <- gen_meta$three_pop

## remove population we have genetic data for but no abundance data
gen <- gen[gen@other$ind.metrics$pop != "WS"]


fst <- pairFST(gen)

fst
```

```
##            TR        BL BR
## TR         NA        NA NA
## BL 0.01437133        NA NA
## BR 0.01358429 0.0156385 NA
```

That is the overall Fst, but we really want to know how that is changing over our three sampling time points. We have three years of data, which happen to span a major rainfall event, from 2006, 2007, and 2008, where there was a major rainfall event in 2007. Let’s look at the pairwise Fst over the three years.

```
fst_by_year <- map(c(2006, 2007, 2008),
                   ~pairFST(gen[gen@other$ind.metrics$year == .x, ]) %>%
                     as.dist() %>%
                     as.matrix())
names(fst_by_year) <- c("2006", "2007", "2008")
fst_by_year
```

```
## $`2006`
##            TR         BR         BL
## TR 0.00000000 0.07423650 0.07492284
## BR 0.07423650 0.00000000 0.04449832
## BL 0.07492284 0.04449832 0.00000000
## 
## $`2007`
##            BL         TR         BR
## BL 0.00000000 0.02405952 0.03053511
## TR 0.02405952 0.00000000 0.02336933
## BR 0.03053511 0.02336933 0.00000000
## 
## $`2008`
##            BL         BR         TR
## BL 0.00000000 0.06184445 0.06037648
## BR 0.06184445 0.00000000 0.04472136
## TR 0.06037648 0.04472136 0.00000000
```

We can visualize that over the three years:

```
fst_df <- imap_dfr(fst_by_year,
                   ~combn(c("BL", "BR", "TR"), 2) %>%
                     t() %>%
                     as.data.frame() %>%
                     rename(pop1 = V1, pop2 = V2) %>%
                     mutate(fst = .x[cbind(pop1, pop2)],
                            year = .y,
                            pop_combo = paste(pop1, pop2, sep = " to ")))

ggplot(fst_df, aes(year, fst)) +
  geom_path(aes(colour = pop_combo, group = pop_combo)) +
  theme_minimal()
```

```
## save for later
write_rds(fst_df, "data/fst_df.rds")
```

So we see that during the rainfall year a precipitous drop in Fst occurs, which has only partially recovered the following year. Now, can we think of a reason why this pattern might occur? We have a hypothesis! First, we’ll lay it out, then we will interrogate its logic using the simulation tools provided by `slimr`. Earlier on we talked about how the desert “turns green” after a big rainfall event. This implies it reverts back to dry or “red” sands afterwards. Generally this is true, however, some patches remain relatively green during the dry periods. These are sometimes know as mesic ‘refugia’. We hypothesized that during dry periods, our mouse population recedes into these separate refugia across the landscape, where they become relatively reproductively isolated. This leads to an inexorable increase in Fst due to a genetic drift without gene flow. Then, when the rain comes, the whole landscape becomes wet, the mice come out to play, spreading throughout, reproducing along the way. The subpopulations of the refugia intermix, leading to an erasure of the genomic differentiation that had been in progress during the dry years. But can this mechanism lead to the level of observed change in such a short period? Are there other possible mechanisms? Let’s explore these ideas with simulation!

#### A simple simulation

To begin setup a simple simulation that captures some of the features of the system we are studying. We will simulate three subpopulations, as we have in our data, and allow migration between them at equal rates. We will include evolutionary processes such as mutation, selection, sexual reproduction, and recombination as additional processes happening in our simulation. Our initial question is simply, can we observe Fst values that change through time in a manner consistent with our observed changes using combinations of just these processes? Thanks to `slimr`, we can continue and write out the logic of our proposed simulation directly in R, using syntax very similar to SLiM (>3.0). We can additionally make the simulation code easy to update with different parameter combinations to try out.

We will use the following parameters in our simulation:

| Parameter | Description |
| --- | --- |
| mut\_rate | mutation rate of simulated population |
| genome\_size | size of simulated genome |
| selection\_strength | mean strength of selection specified as a standard deviation of selection coeffs |
| migration\_rates | rate of migration amongst three pops when abundance is high, must be between 0 and 1 |
| abund\_threshold | abundance (before scaling) above which migration between populations is “turned” on |
| recomb\_rate | recombination rate, a nuisance parameter |
| popsize\_scaling | multiply observed abundances by this value to get total subpop size |

Now for the code:

```
default_genome_size <- 300000

pop_sim <- slim_script(
  
  slim_block(initialize(), {
    
    setSeed(12345);
    initializeMutationRate(slimr_template("mut_rate", 1e-6));
    initializeMutationType("m1", 0.5, "n", 0, slimr_template("selection_strength", 0.1));
    initializeGenomicElementType("g1", m1, 1.0);
    initializeGenomicElement(g1, 0, slimr_template("genome_size", default_genome_size) - 1);
    initializeRecombinationRate(slimr_template("recomb_rate", 1e-8));
    initializeSex("A");
    defineConstant("abund", slimr_inline(pop_abunds, delay = TRUE));
    defineConstant("sample_these", slimr_inline(sample_these, delay = TRUE));
    
  }),
  slim_block(1, {
    
    init_pop = slimr_inline(init_popsize, delay = TRUE)
    
    ## set populations to initial size
    sim.addSubpop("p1", asInteger(init_pop[0]));
    sim.addSubpop("p2", asInteger(init_pop[1]));
    sim.addSubpop("p3", asInteger(init_pop[2]));
    
  }),
  
  slim_block(1, late(), {
    ## get starting population from a file which we will fill-in later
    sim.readFromPopulationFile(slimr_inline(starting_pop, delay = TRUE));
    ## migration on or off flags for pops 1-3 (using tag)
    p1.tag = 0;
    p2.tag = 0;
    p3.tag = 0;
  }),
  
  slim_block(1, 1000, late(), {
    
    ## update generation number
    gen = sim.generation %% 50
    if(gen == 0) {
      gen = 50
    }
    
    ## set population size to observed levels
    p1.setSubpopulationSize(asInteger(ceil(abund[0, gen - 1] * slimr_template("popsize_scaling", 100))));
    p2.setSubpopulationSize(asInteger(ceil(abund[1, gen - 1] * ..popsize_scaling..)));
    p3.setSubpopulationSize(asInteger(ceil(abund[2, gen - 1] * ..popsize_scaling..)));
    
    ## increase migration when above abundance threshold
    if(p1.tag == 0 & abund[0, gen - 1] > slimr_template("abund_threshold", 5)) {
      p2.setMigrationRates(p1, slimr_template("migration_rate", 0))
      p3.setMigrationRates(p1, ..migration_rate..)
      p1.tag = 1;
    } 
    if(p1.tag == 1 & abund[0, gen - 1] <= ..abund_threshold..) {
      p2.setMigrationRates(p1, 0)
      p3.setMigrationRates(p1, 0)
      p1.tag = 0;
    }
    
    if(p2.tag == 0 & abund[1, gen - 1] > ..abund_threshold..) {
      p1.setMigrationRates(p2, ..migration_rate..)
      p3.setMigrationRates(p2, ..migration_rate..)
      p2.tag = 1;
    } 
    if(p2.tag == 1 & abund[1, gen - 1] <= ..abund_threshold..) {
      p1.setMigrationRates(p2, 0)
      p3.setMigrationRates(p2, 0)
      p2.tag = 0;
    }    
    
    if(p3.tag == 0 & abund[2, gen - 1] > ..abund_threshold..) {
      p1.setMigrationRates(p3, ..migration_rate..)
      p2.setMigrationRates(p3, ..migration_rate..)
      p3.tag = 1;
    } 
    if(p3.tag == 1 & abund[2, gen - 1] <= ..abund_threshold..) {
      p1.setMigrationRates(p3, 0)
      p2.setMigrationRates(p3, 0)
      p3.tag = 0;
    }
    
  }),
  
  slim_block(1000, late(), {
    slimr_output_full()
  })
  
)

pop_sim
```

```
## <slimr_script[5]>
## block_init:initialize() {
##     setSeed(12345);
##     initializeMutationRate(..mut_rate..);
##     initializeMutationType("m1", 0.5, "n", 0, ..selection_strength..);
##     initializeGenomicElementType("g1", m1, 1);
##     initializeGenomicElement(g1, 0, ..genome_size.. - 1);
##     initializeRecombinationRate(..recomb_rate..);
##     initializeSex("A");
##     defineConstant("abund", slimr_inline(pop_abunds));
##     defineConstant("sample_these", slimr_inline(sample_these));
## }
## 
## block_2:1 early() {
##     init_pop = slimr_inline(init_popsize);
##     sim.addSubpop("p1", asInteger(init_pop[0]));
##     sim.addSubpop("p2", asInteger(init_pop[1]));
##     sim.addSubpop("p3", asInteger(init_pop[2]));
## }
## 
## block_3:1 late() {
##     sim.readFromPopulationFile(slimr_inline(starting_pop));
##     p1.tag = 0;
##     p2.tag = 0;
##     p3.tag = 0;
## }
## 
## block_4:1:1000 late() {
##     gen = sim.generation%%50;
##     if (gen == 0) {
##         gen = 50;
##     }
##     p1.setSubpopulationSize(asInteger(ceil(abund[0, gen - 1] * ..popsize_scaling..)));
##     p2.setSubpopulationSize(asInteger(ceil(abund[1, gen - 1] * ..popsize_scaling..)));
##     p3.setSubpopulationSize(asInteger(ceil(abund[2, gen - 1] * ..popsize_scaling..)));
##     if (p1.tag == 0 & abund[0, gen - 1] > ..abund_threshold..) {
##         p2.setMigrationRates(p1, ..migration_rate..);
##         p3.setMigrationRates(p1, ..migration_rate..);
##         p1.tag = 1;
##     }
##     if (p1.tag == 1 & abund[0, gen - 1] <= ..abund_threshold..) {
##         p2.setMigrationRates(p1, 0);
##         p3.setMigrationRates(p1, 0);
##         p1.tag = 0;
##     }
##     if (p2.tag == 0 & abund[1, gen - 1] > ..abund_threshold..) {
##         p1.setMigrationRates(p2, ..migration_rate..);
##         p3.setMigrationRates(p2, ..migration_rate..);
##         p2.tag = 1;
##     }
##     if (p2.tag == 1 & abund[1, gen - 1] <= ..abund_threshold..) {
##         p1.setMigrationRates(p2, 0);
##         p3.setMigrationRates(p2, 0);
##         p2.tag = 0;
##     }
##     if (p3.tag == 0 & abund[2, gen - 1] > ..abund_threshold..) {
##         p1.setMigrationRates(p3, ..migration_rate..);
##         p2.setMigrationRates(p3, ..migration_rate..);
##         p3.tag = 1;
##     }
##     if (p3.tag == 1 & abund[2, gen - 1] <= ..abund_threshold..) {
##         p1.setMigrationRates(p3, 0);
##         p2.setMigrationRates(p3, 0);
##         p3.tag = 0;
##     }
## }
## 
## block_5:1000 late() {
##     {sim.outputFull() -> full_output}
## }
## This slimr_script has templating in block(s) block_init
## and block_4 for variables mut_rate and
## selection_strength and genome_size and recomb_rate and
## popsize_scaling and abund_threshold and migration_rate.
```

The above script uses a number of different `slimr` feature that we will explain here. To anyone who has used SLiM before, the above script probably looks pretty familiar. For those of you not familiar with SLiM, we will go over each part of the above script one by one to gain an understanding of what the script is doing. Otherwise, if you’d like to know more about SLiM, I would highly recommend reading the SLiM manual, which is very detailed and packed full of examples.

We start by setting a variable that we will use in the `slimr_script` and later on in the R script as well.

We then use the `slimr_script()` function to create a new `slimr_script` object, which will contain all the information needed to run it in SLiM later. `slimr_script` takes a list of `slimr_block` objects as arguments, which are created with the `slimr_block()` function. `slimr_block` objects contain blocks of SLiM code. Generally, there are two main types of code blocks in SLiM: `initialize` blocks and ‘event’ blocks. Out first block above is an `initialize` block, which sets up the SLiM simulation. All `slimr_script()` calls must contain at least one `initialize` block. Most of the action with `slimr` happens in this block for this script. In it, we set up the simulation with our parameters, using two ‘`slimr` verbs’. ‘`slimr` verbs’ are R functions that can be inserted into SLiM code and which modify the code for various useful purposes. Here we are using `slimr_template()`, which lets you created ‘templated’ variables in a script – these are placemarkers that can be filled in with desired values later. We also use `slimr_inline()`, which let’s you ‘inline’ R objects directly into a script. Inlining means the object and its values are converted into a text representation and passed to SLiM by embedding them directly into the script. This all happens behind the scenes, so you don’t need to worry about it if you are not interested. We have used the special argument `delay = TRUE` to delay the evaluation of `slimr_inline()` until later. This is because we haven’t created the R objects `slimr_inline()` references yet.

The `initialize` block is followed by 4 more `slimr_block` calls, each of which specifies SLiM ‘events’. “block\_2” sets up 3 subpopulations in generation 1, and “block\_3” sets up starting conditions for these subpopulation by reading from a file (which we create later), also in generation 1. “block\_4” contains most of the logic of the simulation, running in every generation from 1 to 1000. Finally, “block\_5” outputs the state of the simulation at the last generation, generation 1000. We assigned the `slimr_script` object to a variable called `pop_sim`, and printed this object, obtaining a pretty output of our script, including special syntax showing our templated variable, which look like this: ..variable\_name.. So, in order to run this script in SLiM, we first need to fill in values for our templated variables, and also make sure we the R objects referred to in `slimr_inline()` calls exist in our R session. We can then “render” our script into a runnable form.

so, how do we fill in the templated variables with values and fill-in our `slimr_inline` calls with actual R objects? For that, we use the `slim_script_render` function. The first thing we need to do before we use it is to create the R objects that `slimr_inline` refers to, otherwise we will get an error about non-existent objects. So what do we need? We need a matrix of abundances for our three subpopulations (for `slimr_inline(pop_abunds)`), a matrix of starting population abundances (for `slimr_inline(init_popsize)`) and a file name pointing to a SLiM population data file containing initial population conditions (for `slimr_inline(starting_pop`). We will later create this file from our `genlight` object using `slimr`. But first, let’s get our objects in order, choose some parameter values and render ourselves a `slimr_script`!

For our population abundances over time, we will use the live-trapping data, and we will create a simplified cycle, taking only part of the sequence, and then repeating it over and over throughout the years. We will sample data between 1995 and 2009, interpolate that to 50 time points, then loop over it inside our `slimr_script`, as we discussed above.

```
pop_abunds <- abund_summ %>%
  filter(date < "2009-10-01" & date > "1995-03-01") %>%
  mutate(three_pop = case_when(pop %in% c("MC", "SS", "WS") ~ "BR",
                               pop %in% c("FRN", "FRS") ~ "BL",
                               pop %in% c("KSE", "CS") ~ "TR")) %>%
  drop_na(three_pop) %>%
  group_by(date, three_pop) %>%
  summarise(abund = mean(abund, na.rm = TRUE),
            .groups = "drop") 

pop_abunds <- pop_abunds %>%
  pivot_wider(names_from = date, values_from = abund) %>%
  as.matrix()

pop_abunds <- rbind(approx(pop_abunds[1, ], n = 50)$y,
                    approx(pop_abunds[2, ], n = 50)$y,
                    approx(pop_abunds[3, ], n = 50)$y)
```

```
## Warning in xy.coords(x, y, setLab = FALSE): NAs introduced by coercion

## Warning in xy.coords(x, y, setLab = FALSE): NAs introduced by coercion

## Warning in xy.coords(x, y, setLab = FALSE): NAs introduced by coercion
```

```
## replace exact zeroes
pop_abunds[pop_abunds == 0] <- 0.02

## set sample times corresponding to our genetic data (roughly)
sample_times <- c("2006" = 40, "2007" = 45, "2008" = 49)

## plot our abundance sequence

plot(pop_abunds[2, ], type = "l", col = "blue")
lines(pop_abunds[1, ], col = "red")
lines(pop_abunds[3, ], col = "green")
abline(v = sample_times)
```

```
## save for later use
write_rds(pop_abunds, "data/pop_abunds.rds")
```

So, that is out `pop_abunds`. For initial population states we are going to base it on data from 2008, which is the third sampling date, and which corresponds to what we think is a panmictic population, which is just after, but not during the big rainfall event. We will get the initial population sizes from here, as well as our starting population data. Our initial population sizes need to be based on this (rather than our abundance data we just generated above), because the number of individuals at the start of our simulation must match the number in the starting population data. The population will then be immediately adjusted to our desired population size by essentially drawing offspring from our smaller starting population pool. This should give us populations with SNP frequencies resembling our actual data at the start of the simulation.

```
## extract 2008 data
gen_2008 <- gen[gen@other$ind.metrics$year == 2008, ]
## count number of individuals in genetic sample per subpopulation
init_popsize <- c(table(pop(gen_2008)))
## set filename to be used for starting pop data (using slim_file to make sure SLiM can find it)
starting_pop = here("sims/starting_pop.txt") %>%
  slim_file()
## setup generations to sample
## we will just sample the three years corresponding to the data, but do it during the last 6 cycles, as "technical" replicates
sample_these <- purrr::map(c(14:19),
                           ~50*.x + sample_times) %>%
  purrr::reduce(union)

## save for later
write_rds(sample_these, "data/sample_these.rds")
write_rds(init_popsize, "data/init_popsize.rds")
```

Okay, now we are ready to use `slim_script_render` to generate a script we can run. The first thing we can try is to render the script without providing a template. Since we have provided defaults for all of our templated variables, this should generate a script with the default values (our ‘default’ script).

```
script_def <- slim_script_render(pop_sim)
```

```
## Warning: Warning: A templated variable was not specified in the template and has been replaced by its default value.
```

```
script_def
```

```
## <slimr_script[5]>
## block_init:initialize() {
##     setSeed(12345);
##     initializeMutationRate(1e-06);
##     initializeMutationType("m1", 0.5, "n", 0, 0.1);
##     initializeGenomicElementType("g1", m1, 1);
##     initializeGenomicElement(g1, 0, 3e+05 - 1);
##     initializeRecombinationRate(1e-08);
##     initializeSex("A");
##     defineConstant("abund", {. = matrix(c(0.926, 0.278, 0.463, 0.557, 0.153, ...), nrow = 3, ncol = 50)});
##     defineConstant("sample_these", {. = c(740, 745, 749, 790, 795, 799, 840, 845, 849, 890, ...)});
## }
## 
## block_2:1 early() {
##     init_pop = {. = c(7, 13, 13)}
##     sim.addSubpop("p1", asInteger(init_pop[0]));
##     sim.addSubpop("p2", asInteger(init_pop[1]));
##     sim.addSubpop("p3", asInteger(init_pop[2]));
## }
## 
## block_3:1 late() {
##     sim.readFromPopulationFile({. = "D:/Projects/slimr_manuscript/sims/starting_pop.txt"});
##     p1.tag = 0;
##     p2.tag = 0;
##     p3.tag = 0;
## }
## 
## block_4:1:1000 late() {
##     gen = sim.generation%%50;
##     if (gen == 0) {
##         gen = 50;
##     }
##     p1.setSubpopulationSize(asInteger(ceil(abund[0, gen - 1] * 100)));
##     p2.setSubpopulationSize(asInteger(ceil(abund[1, gen - 1] * 100)));
##     p3.setSubpopulationSize(asInteger(ceil(abund[2, gen - 1] * 100)));
##     if (p1.tag == 0 & abund[0, gen - 1] > 5) {
##         p2.setMigrationRates(p1, 0);
##         p3.setMigrationRates(p1, 0);
##         p1.tag = 1;
##     }
##     if (p1.tag == 1 & abund[0, gen - 1] <= 5) {
##         p2.setMigrationRates(p1, 0);
##         p3.setMigrationRates(p1, 0);
##         p1.tag = 0;
##     }
##     if (p2.tag == 0 & abund[1, gen - 1] > 5) {
##         p1.setMigrationRates(p2, 0);
##         p3.setMigrationRates(p2, 0);
##         p2.tag = 1;
##     }
##     if (p2.tag == 1 & abund[1, gen - 1] <= 5) {
##         p1.setMigrationRates(p2, 0);
##         p3.setMigrationRates(p2, 0);
##         p2.tag = 0;
##     }
##     if (p3.tag == 0 & abund[2, gen - 1] > 5) {
##         p1.setMigrationRates(p3, 0);
##         p2.setMigrationRates(p3, 0);
##         p3.tag = 1;
##     }
##     if (p3.tag == 1 & abund[2, gen - 1] <= 5) {
##         p1.setMigrationRates(p3, 0);
##         p2.setMigrationRates(p3, 0);
##         p3.tag = 0;
##     }
## }
## 
## block_5:1000 late() {
##     {sim.outputFull() -> full_output}
## }
```

Okay, so now we can see that we have an indication that our R objects will be inserted into the SLiM script. We can try and run this in SLiM now, but we will get an error:

```
test <- slim_run(script_def)
```

```
## 
## 
## Simulation finished with exit status: 1
```

```
## Warning: 
## Failed! Error:
```

```
## ERROR (SLiMSim::InitializePopulationFromFile): initialization file does not exist or is empty.
## 
## Error on script line 26, character 8:
## 
##     sim.readFromPopulationFile("D:/Projects/slimr_manuscript/sims/starting_pop.txt");
##         ^^^^^^^^^^^^^^^^^^^^^^
```

This is because the `starting_pop.txt` doesn’t exist yet, and SLiM has thrown an error telling us this (which slimr passes from SLiM to us through the R console). Let’s create the initial population file using `slim_make_pop_init`. We need to pass this function either a `genlight` or a SNP matrix. Since our `genlight` object has some missing values, and SLiM cannot handle missing values, we will first convert to a SNP matrix, and then fill-in missing values by interpolation. Essentially we just fill-in missing values randomly with a draw from a binomial distribution.

```
snp_mat <- as.matrix(gen_2008)

glimpse(snp_mat)
```

```
##  int [1:33, 1:39978] 0 0 0 0 0 0 0 0 0 0 ...
##  - attr(*, "dimnames")=List of 2
##   ..$ : chr [1:33] "P_herm_281" "P_herm_289" "P_herm_297" "P_herm_265" ...
##   ..$ : chr [1:39978] "15067413-48-G/A" "15067414-5-C/T" "15067416-63-G/A" "15067419-10-G/C" ...
```

```
## replace NAs
nas <- apply(snp_mat, 2, function(x) !any(!is.finite(x)))
snp_mat[ , !nas] <- apply(snp_mat[ , !nas], 2, function(x) {
  tab <- table(x[is.finite(x)]);
  x[!is.finite(x)] <- as.integer(sample(names(tab), sum(!is.finite(x)), replace = TRUE, prob = tab / sum(tab)));
  x
})

glimpse(snp_mat)
```

```
##  int [1:33, 1:39978] 0 0 0 0 0 0 0 0 0 0 ...
##  - attr(*, "dimnames")=List of 2
##   ..$ : chr [1:33] "P_herm_281" "P_herm_289" "P_herm_297" "P_herm_265" ...
##   ..$ : chr [1:39978] "15067413-48-G/A" "15067414-5-C/T" "15067416-63-G/A" "15067419-10-G/C" ...
```

Then, we make our starting population file!

```
sexes <- as.character(gen_2008@other$ind.metrics$Sex)

## sex ratio of sample is skewed to female, so replace actually sexes with extra males
## otherwise simulation crashes because it is unable to sample enough females? Need
## to look more into this bug.
sexes[pop(gen_2008) == "BL"] <- c("M", "F")
```

```
## Warning in sexes[pop(gen_2008) == "BL"] <- c("M", "F"): number of items to
## replace is not a multiple of replacement length
```

```
sexes[pop(gen_2008) == "BR"] <- c("M", "F")
```

```
## Warning in sexes[pop(gen_2008) == "BR"] <- c("M", "F"): number of items to
## replace is not a multiple of replacement length
```

```
sexes[pop(gen_2008) == "TR"] <- c("M", "F")
```

```
## Warning in sexes[pop(gen_2008) == "TR"] <- c("M", "F"): number of items to
## replace is not a multiple of replacement length
```

```
## now make initial population file
slim_make_pop_input(snp_mat, here("sims/starting_pop.txt"), ## filename 
                    sim_gen = 1, ## set generation to first generation
                    ind_pops = gen_2008@other$ind.metrics$three_pop, ## use subpops
                    ind_sex = sexes, ## set sexes
                    ## random positions:
                    mut_pos = sample.int(default_genome_size - 1, nLoc(gen_2008)),
                    version = 3)
```

```
## [1] "D:/Projects/slimr_manuscript/sims/starting_pop.txt"
```

```
## look at the first 50 line to see if it worked:
read_lines(here("sims/starting_pop.txt")) %>%
  head(50)
```

```
##  [1] "#OUT 1 A"                      "Version: 3"                   
##  [3] "Populations:"                  "p1 7 S 0.42857142857142855"   
##  [5] "p2 13 S 0.46153846153846156"   "p3 13 S 0.46153846153846156"  
##  [7] "Mutations:"                    "0 0 m1 233908 0 0.5 p1 1 2"   
##  [9] "1 1 m1 188861 0 0.5 p1 1 8"    "2 2 m1 115563 0 0.5 p1 1 4"   
## [11] "3 3 m1 197484 0 0.5 p1 1 23"   "4 4 m1 38044 0 0.5 p1 1 8"    
## [13] "5 5 m1 240779 0 0.5 p1 1 3"    "6 6 m1 154128 0 0.5 p1 1 4"   
## [15] "7 7 m1 194558 0 0.5 p1 1 5"    "8 8 m1 108248 0 0.5 p1 1 2"   
## [17] "9 9 m1 64572 0 0.5 p1 1 5"     "10 10 m1 207877 0 0.5 p1 1 8" 
## [19] "11 11 m1 243625 0 0.5 p1 1 10" "12 12 m1 278417 0 0.5 p1 1 28"
## [21] "13 13 m1 177657 0 0.5 p1 1 4"  "14 14 m1 276056 0 0.5 p1 1 1" 
## [23] "15 15 m1 283199 0 0.5 p1 1 66" "16 16 m1 2602 0 0.5 p1 1 3"   
## [25] "17 17 m1 139209 0 0.5 p1 1 66" "18 18 m1 13681 0 0.5 p1 1 1"  
## [27] "19 19 m1 21651 0 0.5 p1 1 61"  "20 20 m1 249359 0 0.5 p1 1 12"
## [29] "21 21 m1 98377 0 0.5 p1 1 15"  "22 22 m1 60508 0 0.5 p1 1 62" 
## [31] "23 23 m1 86912 0 0.5 p1 1 8"   "24 24 m1 65026 0 0.5 p1 1 2"  
## [33] "25 25 m1 179118 0 0.5 p1 1 4"  "26 26 m1 172835 0 0.5 p1 1 14"
## [35] "27 27 m1 186257 0 0.5 p1 1 18" "28 28 m1 21785 0 0.5 p1 1 2"  
## [37] "29 29 m1 130856 0 0.5 p1 1 1"  "30 30 m1 153445 0 0.5 p1 1 1" 
## [39] "31 31 m1 111299 0 0.5 p1 1 10" "32 32 m1 115238 0 0.5 p1 1 3" 
## [41] "33 33 m1 87096 0 0.5 p1 1 65"  "34 34 m1 167967 0 0.5 p1 1 1" 
## [43] "35 35 m1 144728 0 0.5 p1 1 1"  "36 36 m1 20188 0 0.5 p1 1 18" 
## [45] "37 37 m1 255815 0 0.5 p1 1 13" "38 38 m1 116350 0 0.5 p1 1 18"
## [47] "39 39 m1 247286 0 0.5 p1 1 3"  "40 40 m1 225973 0 0.5 p1 1 62"
## [49] "41 41 m1 26051 0 0.5 p1 1 11"  "42 42 m1 46943 0 0.5 p1 1 6"
```

And now we can try running our simulation:

```
res <- slim_run(script_def, throw_error = TRUE)
```

```
## Warning: One or more parsing issues, see `problems()` for details
```

```
## 
## 
## Simulation finished with exit status: 0
## 
## Success!
```

Okay, so now we have a working simulation! We’ve currently just output the full population information at the end of the simulation, which doesn’t give us too much information about our question, but just to convince ourselves the simulation produced genetic information, we can extract our output as a `genlight` object and plot it. *Note*: The conversion can take awhile.

```
pop_res <- slimr::slim_extract_genlight(res, "full_output")
## reorder by subpop
pop_res <- pop_res[order(pop(pop_res)), ]
plot(pop_res)
```

So this shows that our three populations are characterized by mostly non-overlapping fixed mutations. This makes sense since our default parameters specified some selection, and no migration. Mutation with positive selection coefficients would have sweeped to fixation, and the lack of migration between subpopulation means that each subpopulation is likely to have had a different set of mutations go to fixation, and no way for these to then spread to other subpopulations.

Okay, to get an idea of what sort of dynamics of Fst we will get with different versions of this simulation, let’s create a version where we calculate Fst during the simulation, and output it for tracking. Then we can make some time-series plots. When it comes to comparing simulation outputs to our actual data we will make yet another version where we will only output just before, during, and just after the population explosions corresponding to rainfall. To add a Fst calculation to our simulation we will use the `slim_function` function, which allows you to specify a SLiM function to use in your SLiM code. This Fst function is based on the one described in the SLiM manual, and can also be found online here: https://github.com/MesserLab/SLiM-Extras/blob/master/functions/calcFST.txt. We have already used the function as an R function when we calculated Fst for our observed data. Now we can convert it to a SLiM function. `slim_function` requires us to specify the function arguments and the function names as character strings, and then the body of the function as `slimr` code. If inserted into a `slim_script` call, the function will then be accessible inside the rest of the code in the `slim_script` call. We can use the previously specified R function code body by simply extracting it using `rlang::fn_body()`. We also need to make sure it is evaluated before insertion, which we assure using the evaluation forcing operator `!!`.

*Note:* The latest version of SLiM (v3.5), which was released after the initial version of this vignette was written, now has a built-in function called `calcFST`, which means we technically no longer need to use a custom function for this functionality. However, we have kept this section of the vignette to show how custom functions can be specified and used in `slimr`.

```
slim_script(
  
  slim_function("o<Subpopulation>$ subpop1", "o<Subpopulation>$ subpop2",
                name = "calculateFST",
                return_type = "f$", body = calculateFST),
  
  slim_block(initialize(), {
    
    setSeed(12345);
    initializeMutationRate(slimr_template("mut_rate", 1e-6));
    initializeMutationType("m1", 0.5, "n", 0, slimr_template("selection_strength", 0.1));
    initializeGenomicElementType("g1", m1, 1.0);
    initializeGenomicElement(g1, 0, slimr_template("genome_size", !!default_genome_size) - 1);
    initializeRecombinationRate(slimr_template("recomb_rate", 1e-8));
    initializeSex("A");
    defineConstant("abund", slimr_inline(pop_abunds, delay = TRUE));
    defineConstant("sample_these", slimr_inline(sample_these, delay = TRUE));
    
  }),
  slim_block(1, {
    
    init_pop = slimr_inline(init_popsize, delay = TRUE)
    
    ## set populations to initial size
    sim.addSubpop("p1", asInteger(init_pop[0]));
    sim.addSubpop("p2", asInteger(init_pop[1]));
    sim.addSubpop("p3", asInteger(init_pop[2]));
    
  }),
  
  slim_block(1, late(), {
    ## get starting population from a file which we will fill-in later
    sim.readFromPopulationFile(slimr_inline(starting_pop, delay = TRUE));
    ## migration on or off flags for pops 1-3 (using tag)
    p1.tag = 0;
    p2.tag = 0;
    p3.tag = 0;
  }),
  
  slim_block(1, 1000, late(), {
    
    ## update generation number
    gen = sim.generation %% 50
    if(gen == 0) {
      gen = 50
    }
    
    ## set population size to observed levels
    p1.setSubpopulationSize(asInteger(ceil(abund[0, gen - 1] * slimr_template("popsize_scaling", 100))));
    p2.setSubpopulationSize(asInteger(ceil(abund[1, gen - 1] * ..popsize_scaling..)));
    p3.setSubpopulationSize(asInteger(ceil(abund[2, gen - 1] * ..popsize_scaling..)));
    
    ## increase migration when above abundance threshold
    if(p1.tag == 0 & abund[0, gen - 1] > slimr_template("abund_threshold", 5)) {
      p2.setMigrationRates(p1, slimr_template("migration_rate", 0))
      p3.setMigrationRates(p1, ..migration_rate..)
      p1.tag = 1;
    } 
    if(p1.tag == 1 & abund[0, gen - 1] <= ..abund_threshold..) {
      p2.setMigrationRates(p1, 0)
      p3.setMigrationRates(p1, 0)
      p1.tag = 0;
    }
    
    if(p2.tag == 0 & abund[1, gen - 1] > ..abund_threshold..) {
      p1.setMigrationRates(p2, ..migration_rate..)
      p3.setMigrationRates(p2, ..migration_rate..)
      p2.tag = 1;
    } 
    if(p2.tag == 1 & abund[1, gen - 1] <= ..abund_threshold..) {
      p1.setMigrationRates(p2, 0)
      p3.setMigrationRates(p2, 0)
      p2.tag = 0;
    }    
    
    if(p3.tag == 0 & abund[2, gen - 1] > ..abund_threshold..) {
      p1.setMigrationRates(p3, ..migration_rate..)
      p2.setMigrationRates(p3, ..migration_rate..)
      p3.tag = 1;
    } 
    if(p3.tag == 1 & abund[2, gen - 1] <= ..abund_threshold..) {
      p1.setMigrationRates(p3, 0)
      p2.setMigrationRates(p3, 0)
      p3.tag = 0;
    }
    
    ## output Fst
    slimr_output(c(calculateFST(p1, p2), calculateFST(p1, p3), calculateFST(p2, p3)), "fsts");
    
  }),
  
  slim_block(1000, late(), {
    sim.simulationFinished()
  })
  
) -> pop_sim_fst
```

Now let’s run that for our defaults to see how it works.

```
fst_script <- slim_script_render(pop_sim_fst)
```

```
## Warning: Warning: A templated variable was not specified in the template and has been replaced by its default value.
```

```
fst_res <- slim_run(fst_script, throw_error = TRUE)
```

```
## 
## 
## Simulation finished with exit status: 0
## 
## Success!
```

So now we can have a look at how the Fst data was returned, which is the default way `slimr` returns data that is a vector:

```
fst_res$output_data
```

```
typeof(fst_res$output_data$data)
```

```
## [1] "character"
```

So it is currently in the form of a character vector separated by spaces. This is pretty straightforward to extract, so let’s do it (*Note*: `slimr` has the ability to automatically convert this output to more user friendly form, but here we will do it the hard way to show how you can flexibly extract different kinds of data, even if a built-in `slimr` function doesn’t exist to handle the type of output you want).

```
extract_fst <- function(fst_res) {
  fst_dat <- fst_res$output_data %>%
        dplyr::select(generation, data) %>%
        dplyr::mutate(data = str_split(data, " ")) %>%
        tidyr::unnest_longer(data, values_to = "fst", indices_to = "index") %>%
        dplyr::mutate(fst = as.numeric(fst), `subpop\npair` = c("BL-BR", "BL-TR", "BR-TR")[index]) 
  fst_dat
}

fst_dat <- extract_fst(fst_res)

ggplot(fst_dat, aes(generation, fst)) +
  geom_path(aes(colour = `subpop\npair`)) +
  scale_y_sqrt() +
  ggforce::facet_zoom(x = generation > 750 & generation < 1000,
                      y = fst > 0.1,
                      horizontal = FALSE,
                      zoom.size = 1)
```

We can see that pairwise Fst rapidly increases between all three subpopulations, reaching nearly its theoretical maximum of 1.0, and then periodically dipping slightly. Let’s look at how this equilibrium pattern corresponds with population sizes. We will look at mean population size and mean Fst.

```
pop_mean <- apply(pop_abunds, 2, mean) %>%
  rep(length.out = n_distinct(fst_dat$generation)) * 100 ## don't forget popsize_scaling

plot_cycle <- function(fst_dat, pop_mean) {
  fst_pop <- fst_dat %>%
    dplyr::group_by(generation) %>%
    dplyr::summarise(fst = mean(fst),
                     .groups = "drop") %>%
    dplyr::mutate(popsize = pop_mean) %>%
    dplyr::filter(generation > 750) ## only take data after equilibrium reached
  
  ggplot(fst_pop, aes(popsize, fst)) +
    geom_path(aes(colour = generation), 
              arrow = arrow(length = unit(0.15, "cm"), type = "closed"), 
              alpha = 0.5) +
    annotate("point", x = fst_pop$popsize[1], fst_pop$fst[1], colour = "red", size = 2) +
    scale_x_log10() +
    scale_colour_viridis_c() +
    theme_minimal()
}

plot_cycle(fst_dat, pop_mean)
```

The plot above shows how Fst and population size are changing through time together for 250 generations (the red dot is the starting point at generation 750). It appears that as population size begins increasing the Fst begins to drop slightly. This drop in Fst continues even as population size begins to decrease again (a lag effect?), then Fst suddenly increases back to nearly 1.0 as population size reaches a low. I suspect this pattern is due to ephemeral effects of increased efficiency of selection when population size is high. As new mutations begin sweeping to fixation there is an intermediate period where they are at intermediate frequency – this has the tendency to lead to reductions in within-population heterozygosity (relative to total heterozygosity), and leads to the subpopulation becoming slightly more similar due to chance collisions between genes being selected in both populations.

Due to overplotting however, some of the details of the plot may be difficult to see. Instead let’s use `gganimate` to create an animated version that might show the dynamics a bit better.

```
animate_cycle <- function(fst_dat, pop_mean) {
  fst_pop <- fst_dat %>%
    dplyr::group_by(generation) %>%
    dplyr::summarise(fst = mean(fst), .groups = "drop") %>%
    dplyr::mutate(popsize = pop_mean) %>%
    dplyr::filter(generation > 250) 
  
  anim <- ggplot(fst_pop, aes(popsize, fst)) +
    geom_point() +
    scale_x_log10() +
    transition_reveal(generation) +
    shadow_wake(0.02, falloff = "sine-in") +
    ggtitle("Generation: {frame_along}") +
    theme_minimal()
  
  animate(anim, nframes = n_distinct(fst_pop$generation) * 5, fps = 30, 
          start_pause = 10, end_pause = 10, renderer = gifski_renderer())
}

animate_cycle(fst_dat, pop_mean)
```

The above animation shows 750 generations of the dynamics, with the black dot showing the position of the population in terms of the mean population size, and the mean pairwise Fst, at different time points. We can clearly see the cycles occurring and its direction.

In any event, this is obviously a pretty unrealistic scenario, and it bears little resemblance to what is happening in our actual data. So, let us try a few other sets of parameters, and see if we can get something closer to what we observe in our data. Now we can finally see how we specify non-default values to our templated variables using `slim_script_render`. Let’s start by changing the `migration_rate` parameter, which controls how much migration we get between subpopulations when they go over their threshold population size. Let’s set it to maximum migration (0.5) – this model is similar to our hypothesis for the pattern we see. We will also set selection to practically zero, so we can isolate drift as a mechanism. We set a parameter value in our script like this:

```
## set migration rate to 0.5 (maximum migration, e.g. 50% of population switches place)
## population will become panmictic over a certain popsize threshold
fst_script_2 <- slim_script_render(pop_sim_fst, template = list(migration_rate = 0.5,
                                                                 selection_strength = 1e-12))
```

```
## Warning: Warning: A templated variable was not specified in the template and has been replaced by its default value.
```

```
## now run the sim and show plots of results
fst_res_2 <- slim_run(fst_script_2, throw_error = TRUE)
```

```
## 
## 
## Simulation finished with exit status: 0
## 
## Success!
```

```
fst_dat_2 <- extract_fst(fst_res_2)

ggplot(fst_dat_2, aes(generation, fst)) +
  geom_path(aes(colour = `subpop\npair`)) +
  scale_y_sqrt() +
  ggforce::facet_zoom(x = generation > 750 & generation < 1000,
                      y = fst < 0.5,
                      horizontal = FALSE,
                      zoom.size = 1)
```

```
plot_cycle(fst_dat_2, pop_mean)
```

```
animate_cycle(fst_dat_2, pop_mean)
```

This shows a very distinct pattern relative to our last run. Now we see that Fst tends to be very low, particularly when population sizes are high (e.g. when migration is “turned on”), as expected. Fst increases in the intervening periods of low population size, which can only be explained by drift. Our Fst values are closer to the magnitude of values we see in our data, which max out at around 0.05, though they still tend to be much higher in this simulation overall.

Let’s try one more scenario before we attempt to more formally compare the simulations to our data. Let’s set our `abund_threshold` controlling at what population size the subpopulation begins exchanging individuals to zero. This means that migration will be on all the time. We will also increase selection.

```
fst_script_3 <- slim_script_render(pop_sim_fst, template = list(migration_rate = 0.5,
                                                                 selection_strength = 0.2,
                                                                 abund_threshold = 0))
```

```
## Warning: Warning: A templated variable was not specified in the template and has been replaced by its default value.
```

```
fst_res_3 <- slim_run(fst_script_3, throw_error = TRUE)
```

```
## 
## 
## Simulation finished with exit status: 0
## 
## Success!
```

```
fst_dat_3 <- extract_fst(fst_res_3)

ggplot(fst_dat_3, aes(generation, fst)) +
  geom_path(aes(colour = `subpop\npair`)) +
  scale_y_sqrt() +
  ggforce::facet_zoom(x = generation > 750 & generation < 1000,
                      y = fst < 0.2,
                      horizontal = FALSE,
                      zoom.size = 1)
```

```
plot_cycle(fst_dat_3, pop_mean)
```

```
animate_cycle(fst_dat_3, pop_mean)
```

We can again see that we get rapid fluctuations in Fst relating to changes in population size. Here they are a bit more rapid but don’t reach the same extremes of Fst. The fluctuations presumably still occur because at very low population size, drift still allows the subpopulations to drift apart due to chance alone, even though they are panmictic (in other words, if you choose any two random subsets of individuals you would expect a higher pairwise Fst during times of low population size). Before we move on, what happens if we reduce the migration rate with an abundance threshold:

```
fst_script_4 <- slim_script_render(pop_sim_fst, template = list(migration_rate = 0.2,
                                                                 selection_strength = 1e-12,
                                                                 abund_threshold = 5))
```

```
## Warning: Warning: A templated variable was not specified in the template and has been replaced by its default value.
```

```
fst_res_4 <- slim_run(fst_script_4, throw_error = TRUE)
```

```
## 
## 
## Simulation finished with exit status: 0
## 
## Success!
```

```
fst_dat_4 <- extract_fst(fst_res_4)

ggplot(fst_dat_4, aes(generation, fst)) +
  geom_path(aes(colour = `subpop\npair`)) +
  scale_y_sqrt() +
  ggforce::facet_zoom(x = generation > 750 & generation < 1000,
                      y = fst < 0.2,
                      horizontal = FALSE,
                      zoom.size = 1)
```

```
plot_cycle(fst_dat_4, pop_mean)
```

```
animate_cycle(fst_dat_4, pop_mean)
```

Again we see those fluctuations. All in all, this suggests that it is likely to be difficult to distinguish what processes are driving the pattern of Fst we see in our study. There clearly are some differences in the details of the dynamics, particularly when you compare our cycle plots between the different scenarios. However, we only have three years of data, just spanning a rainfall (or population explosion) event, which means considerably less information to go on. It looks like it is going to be difficult to disentangle the effects of migration from just the effects of population size, which is perhaps not surprising, given that these two factors are linked in our model (e.g. our hypothesis is that migration increases with high rainfall, which also is linked with population increases). Unfortunately the two processes have a similar effect on Fst, migration creates gene flow which reduces population differentiation, but increasing population increases effective population size, and thus reducing the strength of drift relative to selection, me

So the next thing we should do is get our simulation to output data we can compare directly to our data. In this case we will just output full population information at several time points, and calculate Fst post-hoc using our custom R function to make sure that our methods of calculating Fst are the same. We will also sample the subpopulations in our simulation to try and match sample size with our real data as well.

```
samp_sizes <- table(gen$other$ind.metrics$three_pop, gen$other$ind.metrics$year)

samp_sizes <- do.call(cbind, replicate(6, samp_sizes, simplify = FALSE))

## save for later
write_rds(samp_sizes, "data/samp_sizes.rds")

slim_script(
  
  slim_block(initialize(), {
    
    #setSeed(12345); don't want to set the seed because we want diff each time
    #initializeSLiMOptions(keepPedigrees=T);
    initializeMutationRate(slimr_template("mut_rate", 1e-6));
    initializeMutationType("m1", 0.5, "n", 0, slimr_template("selection_strength", 0.1));
    initializeGenomicElementType("g1", m1, 1.0);
    initializeGenomicElement(g1, 0, slimr_template("genome_size", !!default_genome_size) - 1);
    initializeRecombinationRate(slimr_template("recomb_rate", 1e-8));
    initializeSex("A");
    defineConstant("abund", slimr_inline(pop_abunds, delay = FALSE));
    defineConstant("sample_these", slimr_inline(sample_these, delay = FALSE));
    defineConstant("samp_sizes", slimr_inline(samp_sizes, delay = FALSE));
    
  }),
  slim_block(1, {
    
    init_pop = slimr_inline(init_popsize, delay = TRUE)
    
    ## set populations to initial size
    sim.addSubpop("p1", asInteger(init_pop[0]));
    sim.addSubpop("p2", asInteger(init_pop[1]));
    sim.addSubpop("p3", asInteger(init_pop[2]));
    
  }),
  
  slim_block(1, late(), {
    ## get starting population from a file which we will fill-in later
    sim.readFromPopulationFile(slimr_inline(starting_pop, delay = TRUE));
    ## migration on or off flags for pops 1-3 (using tag)
    p1.tag = 0;
    p2.tag = 0;
    p3.tag = 0;
  }),
  
  slim_block(1, 1000, late(), {
    
    ## update generation number
    gen = sim.generation %% 50
    if(gen == 0) {
      gen = 50
    }
    
    ## set population size to observed levels
    p1.setSubpopulationSize(asInteger(ceil(abund[0, gen - 1] * slimr_template("popsize_scaling", 100))));
    p2.setSubpopulationSize(asInteger(ceil(abund[1, gen - 1] * ..popsize_scaling..)));
    p3.setSubpopulationSize(asInteger(ceil(abund[2, gen - 1] * ..popsize_scaling..)));
    
    ## increase migration when above abundance threshold
    if(p1.tag == 0 & abund[0, gen - 1] > slimr_template("abund_threshold", 5)) {
      p2.setMigrationRates(p1, slimr_template("migration_rate", 0))
      p3.setMigrationRates(p1, ..migration_rate..)
      p1.tag = 1;
    } 
    if(p1.tag == 1 & abund[0, gen - 1] <= ..abund_threshold..) {
      p2.setMigrationRates(p1, 0)
      p3.setMigrationRates(p1, 0)
      p1.tag = 0;
    }
    
    if(p2.tag == 0 & abund[1, gen - 1] > ..abund_threshold..) {
      p1.setMigrationRates(p2, ..migration_rate..)
      p3.setMigrationRates(p2, ..migration_rate..)
      p2.tag = 1;
    } 
    if(p2.tag == 1 & abund[1, gen - 1] <= ..abund_threshold..) {
      p1.setMigrationRates(p2, 0)
      p3.setMigrationRates(p2, 0)
      p2.tag = 0;
    }    
    
    if(p3.tag == 0 & abund[2, gen - 1] > ..abund_threshold..) {
      p1.setMigrationRates(p3, ..migration_rate..)
      p2.setMigrationRates(p3, ..migration_rate..)
      p3.tag = 1;
    } 
    if(p3.tag == 1 & abund[2, gen - 1] <= ..abund_threshold..) {
      p1.setMigrationRates(p3, 0)
      p2.setMigrationRates(p3, 0)
      p3.tag = 0;
    }
    
    ## only run if the generation is in our sample_these list
    if(any(match(sample_these, sim.generation) >= 0)) {
      ## find the sample size that matches the matching "year" for our obs data
      ssizes = drop(samp_sizes[ , which(sample_these == sim.generation)]) 
      ## sample individuals
      ind_sample = c(sample(p1.individuals, ssizes[0]),
                     sample(p2.individuals, ssizes[1]),
                     sample(p3.individuals, ssizes[2]))
      ## output individuals genomes
      slimr_output(ind_sample.genomes.output(), "pop_sample", do_every = 1);
      slimr_output(ind_sample.genomes.individual.subpopulation, "subpops", do_every = 1)
    }
    
  }),
  
  slim_block(1000, late(), {
    sim.simulationFinished()
  })
  
) -> pop_sim_samp
```

Let’s give it a try! Here we will demonstrate another feature of `slimr`. By specifying a `reps` argument to `slim_script_render` we can generate replicate simulations. This returns a `slimr_script_coll` object, which stands for slimr script collection. This can be run using `slim_run` just like a regular `slimr_script`, but each simulation in the collection will be run, either sequentially, or in parallel, depending on what argument we use. We will run in parallel on 6 cores, and run 6 replicates of our entire simulation, with the same parameters. So then we will have 6 replicate samples within each simulation and 6 replicate simulation for a total of 36 replicates for each of our 3 sampling time points.

To run in parallel we must first set a plan for the package `{future}`, which `{slimr}` uses under the hood for parallelism:

```
plan(multisession(workers = 6))
```

Now we can run our script!

```
samp_script <- slim_script_render(pop_sim_samp, template = list(migration_rate = 0.5,
                                                                 selection_strength = 1e-12,
                                                                 abund_threshold = 0),
                                   reps = 6)
```

```
## Warning: Warning: A templated variable was not specified in the template and has been replaced by its default value.
```

```
samp_res <- slim_run(samp_script, throw_error = TRUE,
                     parallel = TRUE)
```

```
## Warning: One or more parsing issues, see `problems()` for details
```

Just to confirm we sampled the generations we wanted:

```
unique(samp_res[[1]]$output_data$generation)
```

```
##  [1] 740 745 749 790 795 799 840 845 849 890 895 899 940 945 949 990 995 999
```

```
sample_these
```

```
##  [1] 740 745 749 790 795 799 840 845 849 890 895 899 940 945 949 990 995 999
```

The last step is to take the output of the simulation, convert it into a set of `genlight` objects, and then calculate Fst on those using the same function used to calculate Fst for our real data. Then we can plot a distribution of expected Fst values at our three time points based on this simulation, and compare that with our real data. We can convert the genomic information to genlight using `slim_extract_genlight()`. By specifying the `name` argument, `slim_extract_genlight()` will only use rows with that name. Additionally we can create a separate genlight for each generation sample by using the `by` argument, like this:

```
slim_genlights <- slim_extract_genlight(samp_res[[1]], name = "pop_sample",
                                       by = "generation")

slim_genlights
```

We now have a tibble containing genlight objects for each generation sample. Let’s plot one to make sure it worked.

```
plot(slim_genlights$genlight[[1]])
```

Now because we output a sample of individuals’ genomes in our simulation, and SLiM doesn’t include information on the subpopulation of origin in its genome output, we also output the associated subpopulation for each genome and included it the `"subpops"` data rows. We can now extract this and reassociate with the genome data. This is possible because SLiM always outputs this data in the same order. Note that we sampled 50 individuals which means we have 100 genomes. These were reassembled into individuals by `slim_extract_genlight()`, and we can likewise aggregate our subpopulation information by noting that SLiM always outputs the two genomes from a single individual one after the other. We can use the `slimr` function `slim_results_to_data()` function which automatically attempts to convert the data stored in our results object to usable data in the form of a tibble. This also works with a

```
slim_subpops <- samp_res[[1]]$output_data %>%
  filter(name == "subpops") %>%
  slim_results_to_data() 

slim_subpops
```

```
slim_subpops <- slim_subpops %>%
  unnest_longer(col = data, values_to = "subpop", indices_to = "genome_num") %>%
  mutate(individual_num = c(rep(1:sum(samp_sizes[ , 1]), each = 2),
                            rep(1:sum(samp_sizes[ , 2]), each = 2),
                            rep(1:sum(samp_sizes[ , 3]), each = 2)) %>%
           rep(6))

slim_subpops
```

```
slim_subpops <- slim_subpops %>%
  group_by(generation, individual_num) %>%
  summarise(subpop = subpop[1],
            .groups = "drop")

slim_subpops
```

Okay, now we can add this to the genlights we generated earlier.

```
## reorder by individual label
slim_genlights$genlight <- map(slim_genlights$genlight,
                               ~.x[order(as.numeric(.x$ind.names)), ])
## add subpop data
slim_genlights$genlight <- map2(slim_genlights$genlight, 
                                slim_subpops %>%
                                  group_split(generation),
                                ~{pop(.x) <- .y$subpop; .x})
```

Let’s look at that plot again but reordered by population.

```
test_gl <- slim_genlights$genlight[[1]]
test_gl <- test_gl[order(pop(test_gl)), ]
plot(test_gl)
```

Now we are ready to calculate our pairwise Fsts for each sample.

```
fst_by_generation_sim <- map(slim_genlights$genlight,
                   ~pairFST(.x) %>%
                     as.dist() %>%
                     as.matrix())
names(fst_by_generation_sim) <- slim_genlights$generation
fst_by_generation_sim
```

```
## $`740`
##                   Subpopulation<p1> Subpopulation<p2> Subpopulation<p3>
## Subpopulation<p1>        0.00000000        0.01502631        0.02842462
## Subpopulation<p2>        0.01502631        0.00000000        0.01465402
## Subpopulation<p3>        0.02842462        0.01465402        0.00000000
## 
## $`745`
##                   Subpopulation<p1> Subpopulation<p2> Subpopulation<p3>
## Subpopulation<p1>       0.000000000        0.01282185       0.009984682
## Subpopulation<p2>       0.012821855        0.00000000       0.009154400
## Subpopulation<p3>       0.009984682        0.00915440       0.000000000
## 
## $`749`
##                   Subpopulation<p1> Subpopulation<p2> Subpopulation<p3>
## Subpopulation<p1>        0.00000000        0.02193978        0.03376151
## Subpopulation<p2>        0.02193978        0.00000000        0.02380561
## Subpopulation<p3>        0.03376151        0.02380561        0.00000000
## 
## $`790`
##                   Subpopulation<p1> Subpopulation<p2> Subpopulation<p3>
## Subpopulation<p1>        0.00000000        0.01434852        0.04054058
## Subpopulation<p2>        0.01434852        0.00000000        0.02042550
## Subpopulation<p3>        0.04054058        0.02042550        0.00000000
## 
## $`795`
##                   Subpopulation<p1> Subpopulation<p2> Subpopulation<p3>
## Subpopulation<p1>       0.000000000       0.007958285       0.009874056
## Subpopulation<p2>       0.007958285       0.000000000       0.006469003
## Subpopulation<p3>       0.009874056       0.006469003       0.000000000
## 
## $`799`
##                   Subpopulation<p1> Subpopulation<p2> Subpopulation<p3>
## Subpopulation<p1>        0.00000000        0.01522112        0.04734702
## Subpopulation<p2>        0.01522112        0.00000000        0.02969295
## Subpopulation<p3>        0.04734702        0.02969295        0.00000000
## 
## $`840`
##                   Subpopulation<p1> Subpopulation<p2> Subpopulation<p3>
## Subpopulation<p1>        0.00000000        0.01176141        0.03527564
## Subpopulation<p2>        0.01176141        0.00000000        0.03276985
## Subpopulation<p3>        0.03527564        0.03276985        0.00000000
## 
## $`845`
##                   Subpopulation<p1> Subpopulation<p2> Subpopulation<p3>
## Subpopulation<p1>       0.000000000       0.012130382       0.009415873
## Subpopulation<p2>       0.012130382       0.000000000       0.009693319
## Subpopulation<p3>       0.009415873       0.009693319       0.000000000
## 
## $`849`
##                   Subpopulation<p1> Subpopulation<p2> Subpopulation<p3>
## Subpopulation<p1>        0.00000000        0.02197344        0.02879841
## Subpopulation<p2>        0.02197344        0.00000000        0.02282966
## Subpopulation<p3>        0.02879841        0.02282966        0.00000000
## 
## $`890`
##                   Subpopulation<p1> Subpopulation<p2> Subpopulation<p3>
## Subpopulation<p1>        0.00000000        0.01886765        0.02638694
## Subpopulation<p2>        0.01886765        0.00000000        0.04743569
## Subpopulation<p3>        0.02638694        0.04743569        0.00000000
## 
## $`895`
##                   Subpopulation<p1> Subpopulation<p2> Subpopulation<p3>
## Subpopulation<p1>       0.000000000        0.01031121       0.008413991
## Subpopulation<p2>       0.010311211        0.00000000       0.007008040
## Subpopulation<p3>       0.008413991        0.00700804       0.000000000
## 
## $`899`
##                   Subpopulation<p1> Subpopulation<p2> Subpopulation<p3>
## Subpopulation<p1>        0.00000000        0.02189298        0.03122043
## Subpopulation<p2>        0.02189298        0.00000000        0.02163579
## Subpopulation<p3>        0.03122043        0.02163579        0.00000000
## 
## $`940`
##                   Subpopulation<p1> Subpopulation<p2> Subpopulation<p3>
## Subpopulation<p1>        0.00000000        0.01895793        0.03215172
## Subpopulation<p2>        0.01895793        0.00000000        0.02103205
## Subpopulation<p3>        0.03215172        0.02103205        0.00000000
## 
## $`945`
##                   Subpopulation<p1> Subpopulation<p2> Subpopulation<p3>
## Subpopulation<p1>       0.000000000       0.010103063       0.005778907
## Subpopulation<p2>       0.010103063       0.000000000       0.008904331
## Subpopulation<p3>       0.005778907       0.008904331       0.000000000
## 
## $`949`
##                   Subpopulation<p1> Subpopulation<p2> Subpopulation<p3>
## Subpopulation<p1>        0.00000000        0.03976261        0.03034681
## Subpopulation<p2>        0.03976261        0.00000000        0.03031384
## Subpopulation<p3>        0.03034681        0.03031384        0.00000000
## 
## $`990`
##                   Subpopulation<p1> Subpopulation<p2> Subpopulation<p3>
## Subpopulation<p1>        0.00000000        0.01290912        0.01893139
## Subpopulation<p2>        0.01290912        0.00000000        0.02755693
## Subpopulation<p3>        0.01893139        0.02755693        0.00000000
## 
## $`995`
##                   Subpopulation<p1> Subpopulation<p2> Subpopulation<p3>
## Subpopulation<p1>       0.000000000        0.01323842       0.008107085
## Subpopulation<p2>       0.013238417        0.00000000       0.018689323
## Subpopulation<p3>       0.008107085        0.01868932       0.000000000
## 
## $`999`
##                   Subpopulation<p1> Subpopulation<p2> Subpopulation<p3>
## Subpopulation<p1>        0.00000000        0.02196998        0.03311335
## Subpopulation<p2>        0.02196998        0.00000000        0.02701812
## Subpopulation<p3>        0.03311335        0.02701812        0.00000000
```

So in this simulation we have very low pairwise fst, including a fair number of negative numbers. This is expected given the high level of migration we used in the simulation (essentially we simulated just one panmictic population, with very weak selection). Now we can write a little function to summarise these results, and plot them along with our observed values. We will also write some new code to get the data from all 6 of our simulation replicates and combine them. Then we can run a few more simulations with different parameters and see how they stack up!

```
sim_result <- samp_res
observed_fst <- fst_df
plot_fst_comparison <- function(sim_result, observed_fst, tech_reps = 6L) {
  slim_genlights <- slim_extract_genlight(sim_result, name = "pop_sample",
                                       by = "generation")
  
  names(sim_result) <- paste0("rep_", seq_along(sim_result))
  
  slim_subpops <- imap_dfr(sim_result, ~ .x$output_data %>%
                            filter(name == "subpops") %>%
                            filter(!duplicated(generation)) %>% ## deals with an intermittent bug
                            slim_results_to_data() %>%
                            unnest_longer(col = data, values_to = "subpop",
                                          indices_to = "genome_num") %>%
                            mutate(individual_num = c(rep(1:sum(samp_sizes[ ,
                                                                            1]), each = 2),
                                                      rep(1:sum(samp_sizes[ , 2]), each = 2),
                                                      rep(1:sum(samp_sizes[ , 3]), each = 2)) %>%
                                     rep(tech_reps),
                                   rep = .y) %>%
                            group_by(generation, individual_num, rep) %>%
                            summarise(subpop = subpop[1],
                                      .groups = "drop"))
  
  ## reorder by individual label
  slim_genlights$genlight <- map(slim_genlights$genlight,
                                 ~.x[order(as.numeric(.x$ind.names)), ])
  ## add subpop data
  slim_genlights$genlight <- map2(slim_genlights$genlight, 
                                  slim_subpops %>%
                                    group_split(rep, generation),
                                  ~{pop(.x) <- .y$subpop; .x})
  
  
  fst_by_year_sim <- map(slim_genlights$genlight,
                     ~pairFST(.x) %>%
                       as.dist() %>%
                       as.matrix())
  years <- rep(c("2006", "2007", "2008"), tech_reps * 6)
  
  
  fst_sim_df <- imap_dfr(fst_by_year_sim,
                     ~combn(c("Subpopulation<p1>", 
                              "Subpopulation<p2>", 
                              "Subpopulation<p3>"), 2) %>%
                       t() %>%
                       as.data.frame() %>%
                       rename(pop1 = V1, pop2 = V2) %>%
                       mutate(fst = .x[cbind(pop1, pop2)],
                              year = years[.y],
                              rep = slim_genlights$rep[.y],
                              generation = slim_genlights$generation[.y],
                              pop_combo = paste(pop1, pop2, sep = " to "))) %>%
    group_by(rep, generation, year) %>%
    summarise(fst = mean(fst), .groups = "drop") %>%
    mutate(iter = rep(1:(tech_reps*6), each = 3))
  
  all_summ <- fst_sim_df %>%
    group_by(year) %>%
    summarise(fst_mean = mean(fst), fst_sd = sd(fst),
              fst = fst_mean,
              .groups = "drop")
  
  fst_obs_df <- observed_fst %>%
    group_by(year) %>%
    summarise(fst_mean = mean(fst),
              fst = fst_mean,
              iter = 99999,
              .groups = "drop")
  
  fst_plot_df1 <- bind_rows(fst_sim_df %>% mutate(type = "simulated"),
                           fst_obs_df %>% mutate(type = "observed")) 
  
  fst_plot_df2 <- bind_rows(all_summ %>% mutate(type = "simulated"),
                           fst_obs_df %>% mutate(type = "observed")) 
  
  p1 <- ggplot(fst_plot_df1, aes(year, fst)) +
    geom_errorbar(aes(ymin = fst_mean - fst_sd,
                      ymax = fst_mean + fst_sd),
                  width = 0.1,
                  data = all_summ) +
    geom_point(aes(colour = type, alpha = type, size = type)) +
    geom_path(aes(colour = type, alpha = type, group = iter)) +
    geom_point(aes(year, fst),  size = 2,
               data = all_summ, inherit.aes = FALSE) +
    geom_path(aes(year, fst), group = 1, data = all_summ, 
              inherit.aes = FALSE) +
    scale_alpha_manual(values = c(1, 0.5)) +
    scale_size_manual(values = c(2, 1)) +
    theme_minimal()
    
  p1
  
}

plot_fst_comparison(samp_res, fst_df)
```

This function plots the means of the pairwise Fst calculated from the simulation along with error bars representing 2 standard deviations around the mean, and the blue lines are each of the 36 replicates (6 temporal times 6 simulation runs). The red dots are the observed data. In this case, the match is not bad, the simulation generates a drop is Fst in 2007, with a rebound in 2008, but overall the Fst values are too low and the rebound is too strong. Let’s try running our simulation again with some of the parameter combinations we explored earlier and see how this plot looks for them!

```
## set migration rate to 0.5 (maximum migration, e.g. 50% of population switches place)
## population will become panmictic over a certain popsize threshold
fst_samp_2 <- slim_script_render(pop_sim_samp, template = list(migration_rate = 0.5,
                                                                selection_strength = 1e-12),
                                  reps = 6)
```

```
## Warning: Warning: A templated variable was not specified in the template and has been replaced by its default value.
```

```
## now run the sim and show plots of results
samp_res_2 <- slim_run(fst_samp_2, throw_error = TRUE,
                       parallel = TRUE)
plot_fst_comparison(samp_res_2, fst_df)
```

Okay, well that simulation produces Fsts within a similar average range as the observed, but interestingly Fst is too high in 2006 and then too low in 2007 and 2008 during and after the rainfall event. This is something that we couldn’t tell from the plots of Fst over time, which showed fluctuations but did not show the detail of exactly when the fluctuations occurred. Let’s try another set of parameters!

```
## set migration rate to 0.5 (maximum migration, e.g. 50% of population switches place)
## population is panmictic all the time with abund_threshold = 0,
## now there is stronger selection too
fst_samp_3 <- slim_script_render(pop_sim_samp, template = list(migration_rate = 0.5,
                                                                selection_strength = 0.2,
                                                                abund_threshold = 0),
                                  reps = 6)
```

```
## Warning: Warning: A templated variable was not specified in the template and has been replaced by its default value.
```

```
## now run the sim and show plots of results
samp_res_3 <- slim_run(fst_samp_3, throw_error = TRUE,
                       parallel = TRUE)
plot_fst_comparison(samp_res_3, fst_df)
```

Now the pattern across years is again similar to the real data, though again the rebound is a little too strong and again overall Fst values are too low. To me, the second simulation that was based most strongly on our hypothesis was quite promising. Let’s tweak the parameters of that one slightly to see if we can get a bit better fit.

```
fst_samp_4 <- slim_script_render(pop_sim_samp, template = list(migration_rate = 0.3,
                                                                selection_strength = 1e-12,
                                                                abund_threshold = 2),
                                  reps = 6)
```

```
## Warning: Warning: A templated variable was not specified in the template and has been replaced by its default value.
```

```
## now run the sim and show plots of results
samp_res_4 <- slim_run(fst_samp_4, throw_error = TRUE,
                       parallel = TRUE)
```

```
## Warning: One or more parsing issues, see `problems()` for details
```

```
plot_fst_comparison(samp_res_4, fst_df)
```

So that is not much better. So far, we haven’t found a perfect simulation, but we have discovered that quite a range of different evolutionary scenarios can create fairly similar patterns. The next step may be to do some formal fitting of the simulation data to the observed using a method such as Approximate Bayesian Computation (ABC). This would be straightforward using `slimr`, since R already has good packages for performing ABC with custom simulation code. It may be the case that for distinguishing the evolutionary forces we are interested in here that we may need more data, or data of a different kind. We could also use simulation to further ask, is there anything we could measure or calculate, besides Fst, from the genomes of individuals, that could better distinguish multiple evolutionary forces? I will leave that project up to future work, but hopefully I have demonstrated the potential of having a tighter integration between data analysis platforms and simulation software.

Last but not least, this is the session information for this run of the above code. We can only guarantee reproducibility of this document if the session where it is being run matches the following:

```
library(sessioninfo)
```

```
## Warning: package 'sessioninfo' was built under R version 4.0.5
```

```
session_info()
```

```
## - Session info ---------------------------------------------------------------
##  setting  value
##  version  R version 4.0.2 (2020-06-22)
##  os       Windows 10 x64 (build 22000)
##  system   x86_64, mingw32
##  ui       RTerm
##  language (EN)
##  collate  English_Australia.1252
##  ctype    English_Australia.1252
##  tz       America/New_York
##  date     2022-03-08
##  pandoc   2.14.0.3 @ C:/Program Files/RStudio/bin/pandoc/ (via rmarkdown)
## 
## - Packages -------------------------------------------------------------------
##  package      * version   date (UTC) lib source
##  ade4         * 1.7-18    2021-09-16 [1] CRAN (R 4.0.5)
##  adegenet     * 2.1.5     2021-10-09 [1] CRAN (R 4.0.5)
##  ape            5.6-2     2022-03-02 [1] CRAN (R 4.0.5)
##  assertthat     0.2.1     2019-03-21 [1] CRAN (R 4.0.2)
##  base64enc      0.1-3     2015-07-28 [1] CRAN (R 4.0.0)
##  bit            4.0.4     2020-08-04 [1] CRAN (R 4.0.2)
##  bit64          4.0.5     2020-08-30 [1] CRAN (R 4.0.2)
##  brio           1.1.3     2021-11-30 [1] CRAN (R 4.0.5)
##  bslib          0.3.1     2021-10-06 [1] CRAN (R 4.0.5)
##  cachem         1.0.6     2021-08-19 [1] CRAN (R 4.0.5)
##  calibrate      1.7.7     2020-06-19 [1] CRAN (R 4.0.2)
##  callr          3.7.0     2021-04-20 [1] CRAN (R 4.0.5)
##  class          7.3-20    2022-01-13 [2] CRAN (R 4.0.5)
##  classInt       0.4-3     2020-04-07 [1] CRAN (R 4.0.2)
##  cli            3.2.0     2022-02-14 [1] CRAN (R 4.0.5)
##  cluster        2.1.2     2021-04-17 [2] CRAN (R 4.0.5)
##  codetools      0.2-18    2020-11-04 [2] CRAN (R 4.0.3)
##  colorspace     2.0-3     2022-02-21 [1] CRAN (R 4.0.5)
##  combinat       0.0-8     2012-10-29 [1] CRAN (R 4.0.0)
##  conflicted   * 1.1.0     2021-11-26 [1] CRAN (R 4.0.5)
##  crayon         1.5.0     2022-02-14 [1] CRAN (R 4.0.5)
##  crosstalk      1.2.0     2021-11-04 [1] CRAN (R 4.0.5)
##  dartR        * 1.9.9.1   2021-05-28 [1] CRAN (R 4.0.5)
##  data.table     1.14.2    2021-09-27 [1] CRAN (R 4.0.5)
##  DBI            1.1.2     2021-12-20 [1] CRAN (R 4.0.5)
##  DEoptimR       1.0-10    2022-01-03 [1] CRAN (R 4.0.5)
##  desc           1.4.0     2021-09-28 [1] CRAN (R 4.0.2)
##  devtools       2.4.3     2021-11-30 [1] CRAN (R 4.0.2)
##  digest         0.6.29    2021-12-01 [1] CRAN (R 4.0.5)
##  directlabels * 2021.1.13 2021-01-16 [1] CRAN (R 4.0.3)
##  dismo          1.3-5     2021-10-11 [1] CRAN (R 4.0.5)
##  doParallel     1.0.17    2022-02-07 [1] CRAN (R 4.0.5)
##  dplyr        * 1.0.8     2022-02-08 [1] CRAN (R 4.0.5)
##  e1071          1.7-9     2021-09-16 [1] CRAN (R 4.0.5)
##  ellipsis       0.3.2     2021-04-29 [1] CRAN (R 4.0.5)
##  evaluate       0.15      2022-02-18 [1] CRAN (R 4.0.5)
##  fansi          1.0.2     2022-01-14 [1] CRAN (R 4.0.5)
##  farver         2.1.0     2021-02-28 [1] CRAN (R 4.0.5)
##  fastmap        1.1.0     2021-01-25 [1] CRAN (R 4.0.2)
##  foreach        1.5.2     2022-02-02 [1] CRAN (R 4.0.5)
##  fs             1.5.2     2021-12-08 [1] CRAN (R 4.0.5)
##  furrr        * 0.2.3     2021-06-25 [1] CRAN (R 4.0.5)
##  future       * 1.24.0    2022-02-19 [1] CRAN (R 4.0.5)
##  gap            1.2.3-1   2021-04-21 [1] CRAN (R 4.0.5)
##  gdata          2.18.0    2017-06-06 [1] CRAN (R 4.0.0)
##  gdistance      1.3-6     2020-06-29 [1] CRAN (R 4.0.2)
##  gdsfmt         1.24.1    2020-06-16 [1] Bioconductor
##  generics       0.1.2     2022-01-31 [1] CRAN (R 4.0.5)
##  genetics       1.3.8.1.3 2021-03-01 [1] CRAN (R 4.0.5)
##  GGally         2.1.2     2021-06-21 [1] CRAN (R 4.0.5)
##  gganimate    * 1.0.7     2020-10-15 [1] CRAN (R 4.0.3)
##  ggforce      * 0.3.3     2021-03-05 [1] CRAN (R 4.0.5)
##  ggplot2      * 3.3.5     2021-06-25 [1] CRAN (R 4.0.5)
##  gifski         1.4.3-1   2021-05-02 [1] CRAN (R 4.0.5)
##  globals        0.14.0    2020-11-22 [1] CRAN (R 4.0.3)
##  glue           1.6.2     2022-02-24 [1] CRAN (R 4.0.5)
##  gridExtra      2.3       2017-09-09 [1] CRAN (R 4.0.2)
##  gtable         0.3.0     2019-03-25 [1] CRAN (R 4.0.2)
##  gtools         3.9.2     2021-06-06 [1] CRAN (R 4.0.5)
##  here         * 1.0.1     2020-12-13 [1] CRAN (R 4.0.2)
##  hierfstat      0.5-10    2021-11-17 [1] CRAN (R 4.0.5)
##  highr          0.9       2021-04-16 [1] CRAN (R 4.0.5)
##  hms            1.1.1     2021-09-26 [1] CRAN (R 4.0.5)
##  htmltools      0.5.2     2021-08-25 [1] CRAN (R 4.0.5)
##  htmlwidgets    1.5.4     2021-09-08 [1] CRAN (R 4.0.5)
##  httpuv         1.6.5     2022-01-05 [1] CRAN (R 4.0.5)
##  igraph         1.2.11    2022-01-04 [1] CRAN (R 4.0.5)
##  iterators      1.0.14    2022-02-05 [1] CRAN (R 4.0.5)
##  jquerylib      0.1.4     2021-04-26 [1] CRAN (R 4.0.2)
##  jsonlite       1.8.0     2022-02-22 [1] CRAN (R 4.0.5)
##  KernSmooth     2.23-20   2021-05-03 [2] CRAN (R 4.0.5)
##  knitr          1.37      2021-12-16 [1] CRAN (R 4.0.5)
##  labeling       0.4.2     2020-10-20 [1] CRAN (R 4.0.3)
##  later          1.3.0     2021-08-18 [1] CRAN (R 4.0.5)
##  lattice        0.20-45   2021-09-22 [2] CRAN (R 4.0.5)
##  leafem         0.1.6     2021-05-24 [1] CRAN (R 4.0.5)
##  leaflet        2.1.0     2022-02-16 [1] CRAN (R 4.0.5)
##  lifecycle      1.0.1     2021-09-24 [1] CRAN (R 4.0.5)
##  listenv        0.8.0     2019-12-05 [1] CRAN (R 4.0.2)
##  lubridate    * 1.8.0     2021-10-07 [1] CRAN (R 4.0.5)
##  magrittr       2.0.2     2022-01-26 [1] CRAN (R 4.0.5)
##  mapview      * 2.10.0    2021-06-05 [1] CRAN (R 4.0.5)
##  MASS           7.3-55    2022-01-13 [2] CRAN (R 4.0.5)
##  Matrix         1.4-0     2021-12-08 [2] CRAN (R 4.0.5)
##  memoise        2.0.1     2021-11-26 [1] CRAN (R 4.0.5)
##  mgcv           1.8-39    2022-02-24 [2] CRAN (R 4.0.5)
##  mime           0.12      2021-09-28 [1] CRAN (R 4.0.5)
##  mmod           1.3.3     2017-04-06 [1] CRAN (R 4.0.2)
##  munsell        0.5.0     2018-06-12 [1] CRAN (R 4.0.2)
##  mvtnorm        1.1-3     2021-10-08 [1] CRAN (R 4.0.5)
##  nlme           3.1-155   2022-01-13 [2] CRAN (R 4.0.5)
##  parallelly     1.30.0    2021-12-17 [1] CRAN (R 4.0.5)
##  patchwork    * 1.1.1     2020-12-17 [1] CRAN (R 4.0.3)
##  pegas          1.1       2021-12-16 [1] CRAN (R 4.0.5)
##  permute        0.9-7     2022-01-27 [1] CRAN (R 4.0.5)
##  pillar         1.7.0     2022-02-01 [1] CRAN (R 4.0.5)
##  pkgbuild       1.3.1     2021-12-20 [1] CRAN (R 4.0.5)
##  pkgconfig      2.0.3     2019-09-22 [1] CRAN (R 4.0.5)
##  pkgload        1.2.4     2021-11-30 [1] CRAN (R 4.0.5)
##  plyr           1.8.6     2020-03-03 [1] CRAN (R 4.0.2)
##  png            0.1-7     2013-12-03 [1] CRAN (R 4.0.0)
##  polyclip       1.10-0    2019-03-14 [1] CRAN (R 4.0.0)
##  PopGenReport   3.0.4     2019-02-04 [1] CRAN (R 4.0.2)
##  prettyunits    1.1.1     2020-01-24 [1] CRAN (R 4.0.2)
##  processx       3.5.2     2021-04-30 [1] CRAN (R 4.0.5)
##  progress       1.2.2     2019-05-16 [1] CRAN (R 4.0.2)
##  promises       1.2.0.1   2021-02-11 [1] CRAN (R 4.0.3)
##  proxy          0.4-26    2021-06-07 [1] CRAN (R 4.0.5)
##  ps             1.6.0     2021-02-28 [1] CRAN (R 4.0.5)
##  purrr        * 0.3.4     2020-04-17 [1] CRAN (R 4.0.2)
##  quadprog       1.5-8     2019-11-20 [1] CRAN (R 4.0.0)
##  R.methodsS3    1.8.1     2020-08-26 [1] CRAN (R 4.0.2)
##  R.oo           1.24.0    2020-08-26 [1] CRAN (R 4.0.2)
##  R.utils        2.11.0    2021-09-26 [1] CRAN (R 4.0.5)
##  R6             2.5.1     2021-08-19 [1] CRAN (R 4.0.5)
##  raster         3.5-15    2022-01-22 [1] CRAN (R 4.0.5)
##  RColorBrewer   1.1-2     2014-12-07 [1] CRAN (R 4.0.0)
##  Rcpp           1.0.8     2022-01-13 [1] CRAN (R 4.0.5)
##  readr        * 2.1.2     2022-01-30 [1] CRAN (R 4.0.5)
##  remotes        2.4.2     2021-11-30 [1] CRAN (R 4.0.5)
##  reshape        0.8.8     2018-10-23 [1] CRAN (R 4.0.2)
##  reshape2       1.4.4     2020-04-09 [1] CRAN (R 4.0.2)
##  rgdal          1.5-28    2021-12-15 [1] CRAN (R 4.0.5)
##  RgoogleMaps    1.4.5.3   2020-02-12 [1] CRAN (R 4.0.2)
##  rlang          1.0.2     2022-03-04 [1] CRAN (R 4.0.5)
##  rmarkdown      2.12      2022-03-02 [1] CRAN (R 4.0.5)
##  robustbase     0.93-9    2021-09-27 [1] CRAN (R 4.0.5)
##  rprojroot      2.0.2     2020-11-15 [1] CRAN (R 4.0.3)
##  rstudioapi     0.13      2020-11-12 [1] CRAN (R 4.0.3)
##  sass           0.4.0     2021-05-12 [1] CRAN (R 4.0.5)
##  satellite      1.0.4     2021-10-12 [1] CRAN (R 4.0.5)
##  scales         1.1.1     2020-05-11 [1] CRAN (R 4.0.2)
##  seqinr         4.2-8     2021-06-09 [1] CRAN (R 4.0.5)
##  sessioninfo  * 1.2.2     2021-12-06 [1] CRAN (R 4.0.5)
##  sf           * 1.0-6     2022-02-04 [1] CRAN (R 4.0.5)
##  shiny          1.7.1     2021-10-02 [1] CRAN (R 4.0.5)
##  slimr        * 0.1.0     2022-03-07 [1] local
##  SNPRelate      1.22.0    2020-04-27 [1] Bioconductor
##  sp             1.4-6     2021-11-14 [1] CRAN (R 4.0.5)
##  StAMPP         1.6.3     2021-08-08 [1] CRAN (R 4.0.5)
##  stringi        1.7.6     2021-11-29 [1] CRAN (R 4.0.5)
##  stringr      * 1.4.0     2019-02-10 [1] CRAN (R 4.0.2)
##  terra          1.5-21    2022-02-17 [1] CRAN (R 4.0.5)
##  testthat       3.1.2     2022-01-20 [1] CRAN (R 4.0.5)
##  tibble       * 3.1.6     2021-11-07 [1] CRAN (R 4.0.5)
##  tidyr        * 1.2.0     2022-02-01 [1] CRAN (R 4.0.5)
##  tidyselect     1.1.2     2022-02-21 [1] CRAN (R 4.0.5)
##  tweenr         1.0.2     2021-03-23 [1] CRAN (R 4.0.5)
##  tzdb           0.2.0     2021-10-27 [1] CRAN (R 4.0.5)
##  units          0.8-0     2022-02-05 [1] CRAN (R 4.0.5)
##  usethis        2.1.5     2021-12-09 [1] CRAN (R 4.0.5)
##  utf8           1.2.2     2021-07-24 [1] CRAN (R 4.0.5)
##  vctrs          0.3.8     2021-04-29 [1] CRAN (R 4.0.5)
##  vegan          2.5-7     2020-11-28 [1] CRAN (R 4.0.3)
##  vroom          1.5.7     2021-11-30 [1] CRAN (R 4.0.5)
##  webshot        0.5.2     2019-11-22 [1] CRAN (R 4.0.2)
##  withr          2.5.0     2022-03-03 [1] CRAN (R 4.0.5)
##  xfun           0.30      2022-03-02 [1] CRAN (R 4.0.5)
##  xtable         1.8-4     2019-04-21 [1] CRAN (R 4.0.2)
##  yaml           2.3.5     2022-02-21 [1] CRAN (R 4.0.5)
##  zeallot        0.1.0     2018-01-28 [1] CRAN (R 4.0.2)
## 
##  [1] C:/Users/rdinn/Documents/R/win-library/4.0
##  [2] C:/R/R-4.0.2/library
## 
## ------------------------------------------------------------------------------
```
